# Vineyard footprint revealed by honey bees used as sentinels for organic and inorganic contaminants in contrasted environments

**DOI:** 10.64898/2025.12.16.694429

**Authors:** Léa Tison, Céline Franc, Louisiane Burkart, Camille Larrue, Camille Grosgeorge, Thierry Dalix, Adrien Rusch, Gaëtane Le Provost, Gilles de Revel, Denis Thiéry

## Abstract

The contamination of non-target organisms by elements and pesticides can adversely affect biodiversity and key ecosystem services such as pollination. In this study, we used honey bee colonies as sentinels of environmental contamination in four distinct habitats - urban, suburban, forest, and vineyard - and collected bees every three weeks from May to November. Pesticide residues were extracted using an adapted QuEChERS method and analyzed by LC-MS/MS, while inorganic major and trace elements were measured by ICP-MS and ICP-AES. Our results show that land-use and seasonal changes shape contaminant profiles in bees. Bees from vineyards exhibited the highest contaminant diversity and loads, indicating a particularly intense contamination profile in this environment. This reflects exposure to multiple pesticide inputs - such as copper - which can accumulate over time to potentially harmful levels. Pollinators like honey bees provide effective indicators of environmental risk, as their contaminant loads may affect honey bee health and can reveal broader ecological impacts.

## Introduction

Worldwide increase in pesticide use raises significant concerns and has become a serious global issue with side effects outweighing expected benefits (Scheringer & Schulz, 2025). A growing body of evidence shows that pesticide use leads to widespread contamination of all environmental compartments (Tang et al., 2021; Zaller et al., 2023; Rigal & Perrot, 2025). The contamination of non-target organisms and ecosystems by pesticides residues is closely linked to non-negligible risks for overall biodiversity and associated ecosystem services such as pollination (Beketov et al., 2013; Rigal et al., 2023; Tison et al., 2024; Wan et al., 2025; Stemmelen et al., 2025). Pesticides residues are not the only contaminants present in our environment. Human activities have raised environmental levels of major and trace metals elements far above the natural baseline (Ondrasek et al., 2025) up to potentially toxic levels for non-targeted fauna.

Such as with pesticides residues, bees seem unable to detect field-realistic concentrations of elements, exposing them chronically (Raine & Gill, 2015; Monchanin et al., 2022). Hence, several elements have been repeatedly detected in foraging bees, with levels correlating strongly with ambient environmental contamination (Perugini et al., 2011; Lambert et al., 2012; Ruschioni et al., 2013; Mair et al., 2023). Honey bees can thus serve to detect temporal and spatial patterns in environmental concentrations of contaminants, even at relatively low levels (van der Steen et al., 2012), and *Apis mellifera* has emerged as a reliable biosentinel across diverse environments, due to its extensive foraging range (3 km in average), large diet breadth, and its ability to store food in the hive (Giglio et al., 2017; Cunningham, Crane, et al., 2022). Consequently, honey bees are more and more used to assess the risks of environmental contamination on biodiversity and ecosystem services (Badiou-Bénéteau et al., 2013; Gutiérrez et al., 2015; Girotti et al., 2020). Foraging bees can encounter potentially harmful chemicals in airborne dusts and clouds (Thimmegowda et al., 2020; Bianco et al., 2025), plant nectar and pollen (Perugini et al., 2011; Xun et al., 2018; Cappellari et al., 2024), and water (Li et al., 2016; Lorenz et al., 2025). All bees from a colony can ultimately ingest contaminants via the consumption of pollen and honey, or while in contact with nest materials (Cozmuta et al., 2012; Calatayud-Vernich et al., 2018; Glinski et al., 2024).

They face multiple chemical stress from both pesticides and elements with direct lethal toxicity but also a range of sub-lethal effects that impair sneakily bee health and colony function. Their toxic effects range from altered metabolism and immune system disruptions (Perugini et al., 2011; Lambert et al., 2012; Di Prisco et al., 2013; Pamminger et al., 2018) to developmental and cognitive deficits such as the impairment of learning, memory, and navigation that reduce foraging effectiveness and fitness (Tison et al., 2016; Burden et al., 2016; Siviter et al., 2018; Monchanin et al., 2021). Recent reviews have highlighted the complexity of pesticide mixture toxicity and the need to jointly assess pesticide and trace metal exposures in pollinators, particularly within EU risk assessment frameworks addressing cumulative stressors(EFSA Scientific Committee et al., 2021). Since pesticide risk is determined by hazard and exposure, it is crucial to assess how non-target organisms interact with the presence and type of chemical products (Stanley et al., 2015; Sponsler et al., 2019). The exposure of bees to both elements and pesticides residues is seldom investigated, despite bees frequently and simultaneously encountering these contaminants. While many studies focused on the impact of one class of contaminant, very few have combined multiple contaminants within a single sentinel framework along an anthropization gradient, ranging from unmanaged habitats to highly disturbed agricultural habitats like vineyards.

Vineyard-dominated landscapes are characterized by the intensive use of pesticides, including synthetic organic substances as well as inorganic fungicides such as the major elements copper and sulfur. Copper, which is authorized in organic viticulture despite accumulation concerns and regulatory restrictions in some EU countries (e.g., Netherlands, Denmark), exemplifies this challenge (European Food Safety Authority (EFSA) et al., 2018). Pesticide spraying is not confined to vineyard plots, since spray drift and surface run-off can extend contamination to adjacent habitats, affecting soils, water bodies, and non-target organisms (Vischetti et al., 2008; Albaseer et al., 2025). As a result, vineyard agroecosystems often represent hotspots of exposure to contaminants, generating a substantial ecotoxicological burden for non-targeted species with potentially significant impacts on local biodiversity and ecosystem functioning (Stemmelen et al., 2025). In this context, pollinators such as honey bees offer a valuable lens for assessing environmental risks, as their health and exposure patterns can provide early-warning signals of broader ecological impacts (Cunningham, Tran, et al., 2022).

In this study, we deployed honey bee colonies as environmental sentinels across four contrasted environments with distinct ecological characteristics: urban areas, suburban areas, forested areas and vineyards. Our objectives were to (i) assess how different environmental contexts leave detectable ‘chemical footprints’ (diversity and loads of contaminants) on honey bees serving here as biosentinels, and (ii) identify specific environments, substances and concentration levels that may pose risks to honey bees. We hypothesized that (i) environments with intensive chemical use (e.g., vineyards) leave a stronger ecological footprint on honey bees compared to habitats with low chemical input (e.g., forests); (ii) vineyards leave a distinct footprint on honey bees, reflecting combined exposure to both historical and current inputs of organic and inorganic pesticides; (iii) some substances accumulate in living organisms over time, leading to potential bioaccumulation at levels that may become harmful.

## Material and methods

### Study sites

In this study, we considered four sites. All sites are located in Gironde, in southwestern France: one is located close to INRAE Grande Ferrade research center, Villenave d’Ornon, one is located close to the city of Gujan-Mestras, another one is located close to the “Lande de Mejos”, and the last one is located close to the village of Saint-Magne de Castillon. Land use around the four study sites was characterized within a 3 km radius of the monitored colonies (Figure 1) using QGIS Geographic Information System, complemented with Google satellite images and OSO Land Cover Map 2020 (Thierion et al., 2022). To characterize each site, we considered the following categories: urban areas (including residential areas, roads, and industrial zones), grassland, vineyard, forest, water bodies, and annual crops (i.e., oilseed and corn). The proportion of each land-use type was calculated within the buffer and this information was used for the a priori classification of the different sites: Urban ( > 75 % urban fabric), Suburban (> 50 % urban fabric), Forest (> 75 % forest) and Vineyard (> 50 % vineyards).

**Figure 1.**
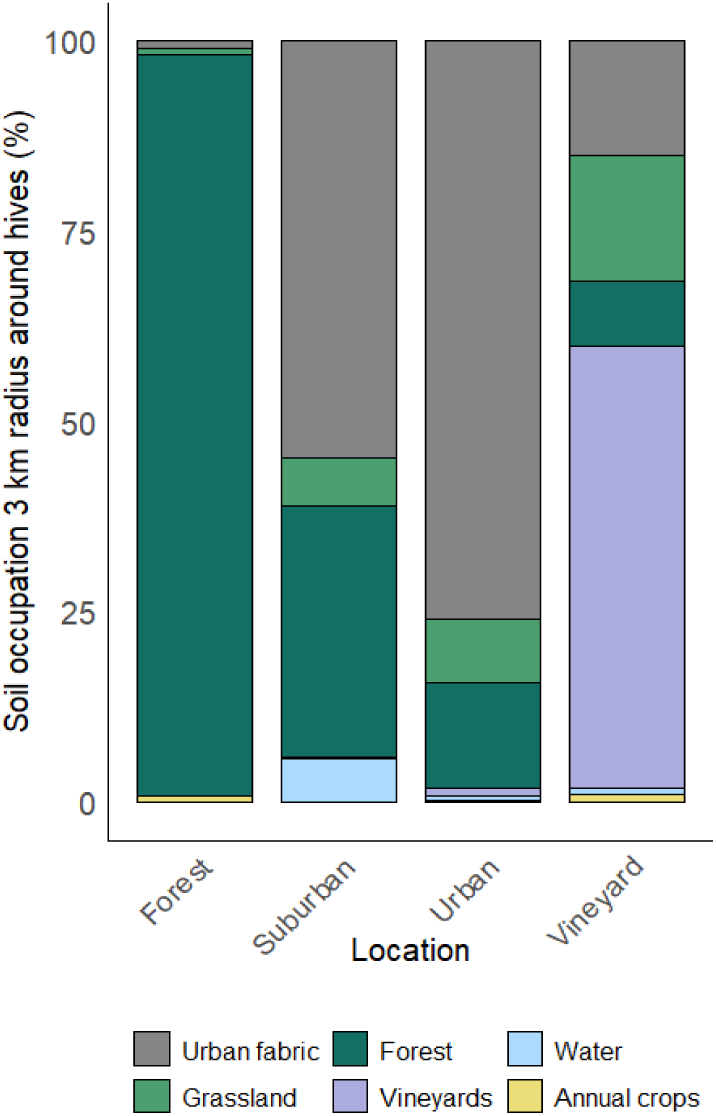
Land use in a 3 km radius around hives installed in four contrasted locations.

### Sample collection

Eight honey bee colonies (*A.mellifera*) were placed in the four different environments (two colonies per site). Returning honey bees foragers were sampled at the entrance of the hive every three weeks between May 3rd and October 20th (9 samplings). Honey bees usually live around 5 to 6 weeks in spring and summer but become foragers only at the end of their lives for 7 to 12 days (Rueppell et al., 2007). Collected honey bees were kept at 0°C immediately after sampling and subsequently stored at -20 °C until analysis at the laboratory. For the analysis of pesticide residues, 10 honey bees were collected per colony, sampling and site. For setting the calibration curves and recovery rates additional honey bees were sampled in a colony located in an uncontaminated and previously sampled site in Bouliac, Gironde. For the analysis of elements, about 5 individuals were collected per colony, sampling and site. In order to characterize the overall contamination profile (contact, ingestion) which reflects real-world exposure conditions, bees were not washed prior to chemical analysis.

### Analytes and reagents

Details of molecules used in the study and related analytical standards are given in Supporting Information Table S1. Triphenylphosphate (TPP) was used as generic internal standard and all molecules were purchased from Sigma-Aldrich Chimie (St Quentin Fallavier, France). Isotopically labelled internal standards carbendazim-d_4_ (99.3 % isotopic purity), fludioxonil-^13^C_2_ (99.6 % isotopic purity) tebufenozide-d_9_ (98.9 % isotopic purity) and, tebuconazole-d_6_ (100 ng.μL^-1^ in acetone, 97.5 % isotopic purity) were supplied by Cil Cluzeau Info Labo (Sainte-Foy-La-Grande, France). All pesticide standards were in the range 97 - 99.9 % chemical purity. LC-MS grade acetonitrile (ACN LC-MS, ≥ 99.9 % purity) was obtained from VWR International SAS (Rosny-sous-Bois, France) and glacial acetic acid from Sigma Aldrich (same source as above). Concentrations of the internal standard working mixture were: TPP 500 ng.mL^-1^, carbendazim-d_4_ 545 ng.mL^-1^, fludioxonil-^13^C_2_ 1 µg.mL^-1^, tebufenozide-d_9_ 10 µg.mL^-1^ and, tebuconazole-d_6_ 550 ng.mL^-1^ in ACN LC-MS. The stock solutions of the pesticide mixtures used for calibration, evaluation of recoveries and precisions were prepared in ACN LC-MS at a concentration close to 20 mg.L^-1^. Further dilutions were applied to obtain solutions in ACN LC-MS and acetic acid mixture (99:1) at 40, 20, 10, 4, 1, and 0.2 µg.L^-1^ equivalent to 200, 100, 50, 20, 5, and 1 ng.g^-1^ concentrations in bees. All solutions were stored at −20 °C in the dark. Anhydrous sodium acetate (NaOAc, > 99%), magnesium sulfate (MgSO_4,_ ≥99.5%), and zirconium dioxide-based sorbent (Supel QuE Z-Sep+/MgSO_4_ 2 mL centrifuge tubes) were supplied by Sigma-Aldrich (Saint-Quentin-Fallavier, France). Water was purified through a GENPURE UV-TOC system by Thermo Fischer Scientific (Illkirch-Graffenstaden, France). For the analysis of elements, all reagents were of suprapur grade or equivalent: Nitric acid (HNO₃, 65%, Sharlau, France), Hydrofluoric acid (HF, 40%, Suprapur, Merck, Germany), Hydrogen peroxide (H₂O₂, 30%, Baker, USA), Ultrapure water (18.2 MΩ·cm, Milli-Q, Millipore, France).

### Pesticide residues extraction and analysis

To meet the minimum sample weight required for sensitive pesticide residue analysis but with the limitation of masking within-colony variablilty, pools of 10 foragers of *A. mellifera* per colony were created. The bees were put in liquid nitrogen before grinding in stainless steel 10 mL-grinding jars (cat. no. 69985 from Qiagen, Hilden, Germany) for 6 x 30 seconds at a frequency of 30 Hz (1800 oscillations per minute) using the crusher TissueLyser II (Qiagen, Hilden, Germany). The mash was homogenized, and 250 ± 1 mg of sample was transferred into a 15 mL Falcon tube. For each colony, this was replicated 3 times (triplicate). Samples were then extracted using the QuEChERS method described in Tison et al. (2023) and HPLC-MS/MS analysis was performed using the same instrument and conditions. The retention times, repeatability (Relative Standard Deviation), recovery, and the Limits of Detection and Quantification (LOD/LOQ) of each of the 57 analytes are listed in Supporting Information Table S1.

### Mineralization of elements

Before mineralization, approximately 300 mg of whole bees were weighed using a Radwag analytical balance (precision 0.1 mg) into 50 mL PTFE tubes. Samples were then dried in an oven at 103°C to determine moisture content prior to digestion. Mineralization was carried out using an Environmental Express automated digestion system (Charleston, USA). For each tube, homogenized samples were treated with a mixture of HNO₃, HF, and H₂O₂ according to a controlled digestion program. After complete evaporation, digests were brought to volume with ultrapure water containing 5 % nitric acid (v/v). Reagent blanks and control samples were included for quality control.

### Quantification by ICP-AES and ICP-MS

The elements Aluminium, Calcium, Copper, Iron, Sulfur, Magnesium, Manganese, Phosphorous, Potassium and Zinc were determined by ICP-AES (Varian 725-ES, Varian, Palo Alto, USA) equipped with a V-Groove nebulizer. Wave lengths for each element is given in Supporting Information (Table S2.A). The elements Arsenic, Cadmium, Cobalt, Chromium, Molybdenum and Lead were quantified by ICP-MS (Agilent 7850, Agilent Technologies, Santa Clara, USA) equipped with a Scott-type spray chamber and a Micromist nebulizer. Instrumental conditions were optimized daily using multi-element standard solutions. Rhodium was used as internal standard for all elements in ICP-MS to correct for matrix effects. Isotopes are given in Supporting Information (Table S2.B). Recovery rates were between 96.1 and 106.5 % and are given for each analyzed element in Table S2. Calibrations were performed using single-element standard solutions prepared by dilution of certified stock solutions. An external linear calibration was applied, with systematic verification of blanks and accuracy checks using certified reference materials (V 463 whole plant, NIST SRM 1547 – Peach Leaves, used for cross-validation).

### Statistical analysis

Principal Component Analysis (PCA) was performed on scaled land-use variables to explore patterns of contaminant exposure across study sites (Urban, Suburban, Forest, Vineyard) with the ‘FactoMineR’ package. Group centroids and variable loadings were visualized using biplots to interpret spatial and environmental gradients. Linear mixed-effect models were fitted to compare element concentrations across the four study sites using the packages ’lme4’ (Bates et al., 2015) and ’lmerTest’ (Kuznetsova et al., 2017). With only two colonies per site, statistical power is limited, although repeated measures across dates partially compensate for this constraint. In each model, Element and Location were included in interaction as fixed effects, and both Date and Hive were included as random effects to account for seasonality in element concentration variability and for the repeated measures structure of the design, respectively. Element concentrations measured in honey bees across the four locations were log-transformed to improve normality. Model assumptions of normality (Q-Q plots), homoscedasticity (residual plots vs. fitted values), and absence of substantial serial correlation were validated. For each element, six planned contrasts were tested using the ’emmeans’ package. No correction for multiple comparisons was applied, as these were pre-specified contrasts analyzed independently for each element. Results were visualized with faceted plots showing estimated differences and 95% confidence intervals, highlighting statistically significant site differences (p < 0.05). We then focused on the substances found at higher levels in the Vineyard location in order to evaluate more precisely their accumulation in honey bees. Concentrations of Cu, Co, Cr, S, As and Al were analysed using individual hive observations. Mixed-effect models with month as a fixed covariate and Hive as a random intercept were fitted for each element to evaluate concentration trends over time, allowing slope estimates and associated p-values to be retrieved while accounting for the repeated measures structure within hives. Pesticide treatments applied in the nearest plot obtained from the wine-grower were visually added to the regression plots. Pesticide residues data were log-transformed (log 10 conc +1) in order to generate a heatmap of maximum pesticide residue concentrations measured in honey bees across the four study sites. Fungicide richness (number of distinct fungicides detected per sample) was compared across locations using a Friedman test followed by post-hoc pairwise comparisons with Bonferroni correction. Temporal trends in fungicide richness within each location were assessed using Spearman rank correlations. For calculations of means (Table S3) and graphical representations, pesticide residue concentrations above the limits of detection (≥LOD) but below the limits of quantification (<LOQ) were assigned the LOD value. All statistics and figures were made using R version 4.5.0 (R Core Team. (2025)).

## Results

### Major and trace elements

Elements concentrations varied meaningfully among the four sampling locations, with specific elements linked to distinct land-use or environmental types (Fig. 1 and 2). The PCA biplot (Fig. 2) displays the distribution of sampling sites categorized into our four location groups—Urban, Suburban, Forest, and Vineyard—based on element composition. The four groups are visually well separated along the first two principal components, with distinct clusters evident for each location, as indicated by the colored ellipses. Trace elements such as Lead (Pb), Cadmium (Cd) and Molybdenum (Mo) are primarily associated with Urban and Suburban locations. The element Mn (Manganese) is strongly aligned with the Forest location. Elements such as Arsenic (As), Cobalt (Co), Zinc (Zn), Aluminum (Al), Iron (Fe), Chromium (Cr), and Copper (Cu) form a cluster directing towards the Vineyard location.

**Figure 2.**
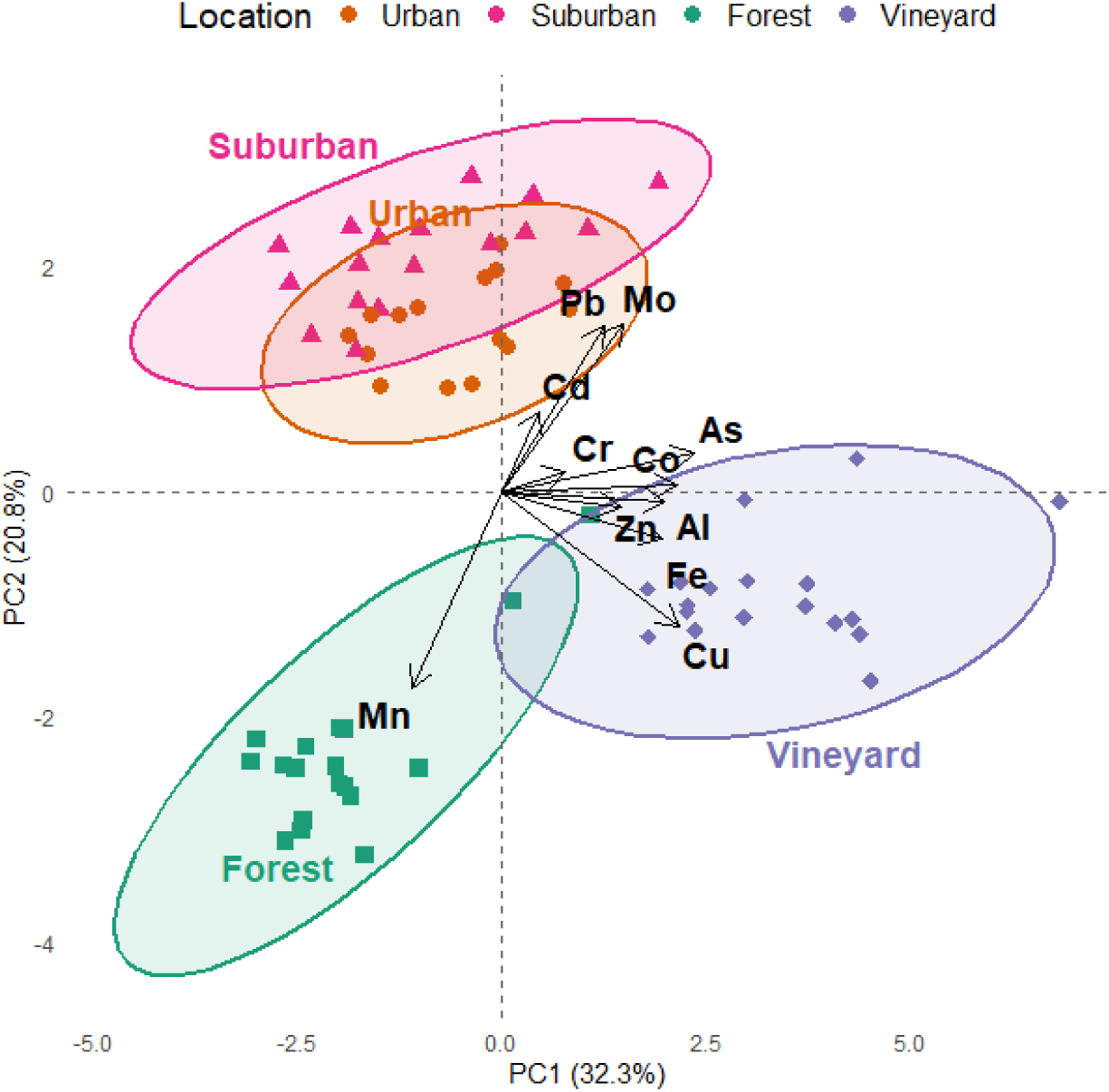
Principal Component Analysis (PCA) biplot of bee elements. Colored points correspond to sampling locations (Forest, Suburban, Urban, or Vineyard). Ellipses indicate 95% confidence intervals for each location group. Black arrows represent the projection of elements with arrow direction and length reflecting their contribution to site distribution along the principal components. PC1 and PC2 axes display the main variance directions in the dataset (32.3 % and 20.8 % respectively, PC3 accounted for 11.5% of the total variance).

The upper panel (A) of Figure 3 presents the estimated effect of the vineyard location compared to the three other locations (Forest, Suburban, and Urban) for each element concentrations based on the fitted linear mixed-effects models. The lower panel (B) show the comparison of element concentrations between the other locations (Forest, Suburban, and Urban). These results highlight the strong across-locations variability in the concentration of chemical elements found in bees, with Vineyard and occasionally Urban sites standing out for higher contaminant loads.

**Figure 3.**
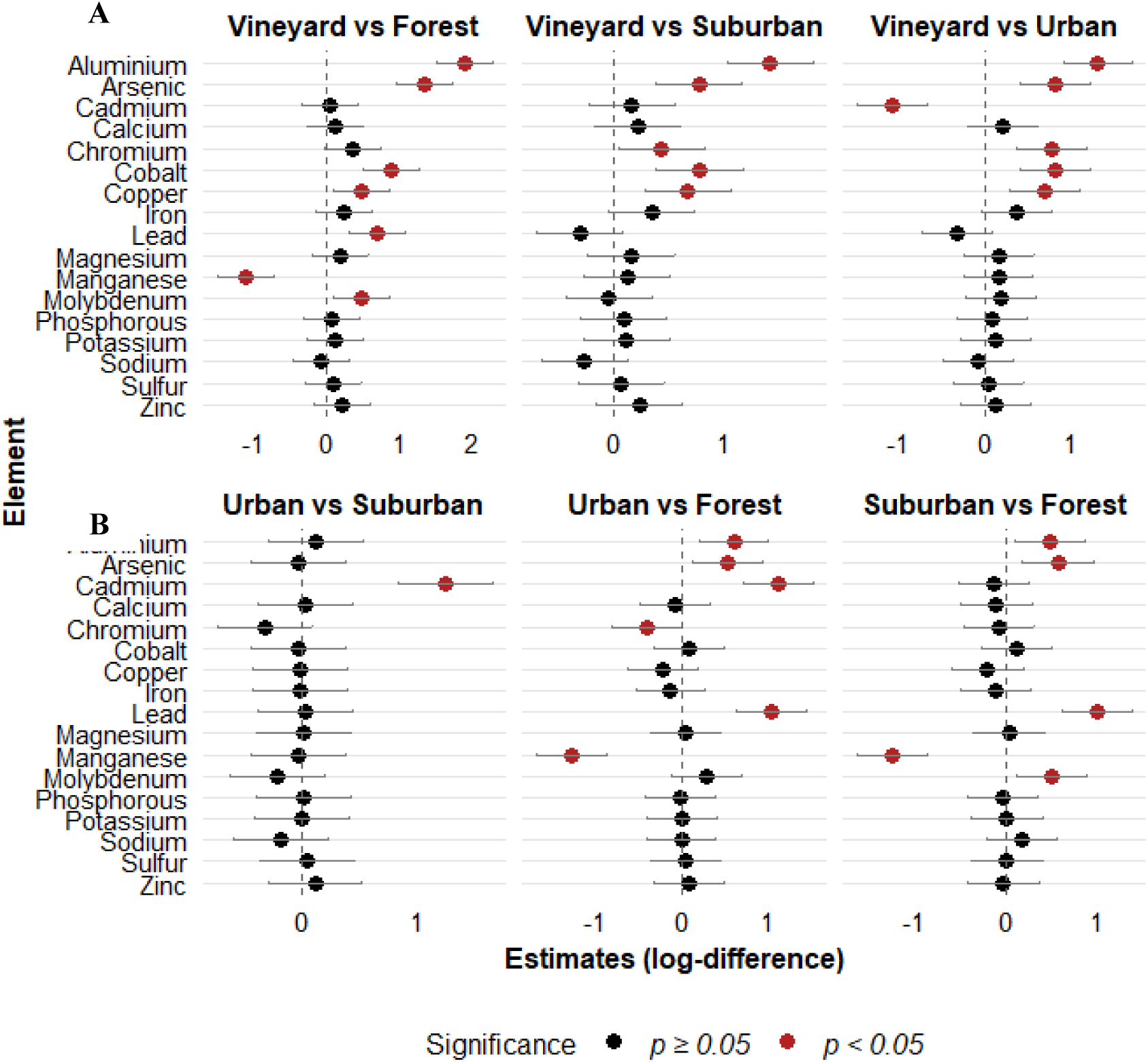
Effects of environment types on element concentrations found in honey bees. The figure presents post-hoc pairwise comparisons of estimated marginal means between locations for each element using Tukey adjustment via the ‘emmeans’ package. Each panel corresponds to one of the contrasts. Metals are displayed on the vertical axis, while the horizontal axis shows the estimated difference in log-concentration between locations. Significant differences (p < 0.05) are shown in red. Horizontal error bars represent 95% confidence intervals of the estimates.

Key findings indicate significantly higher concentrations of several metals at the Vineyard location compared the other sites (Fig. 4A), including Al, As, Cr (except for Vineyard vs Forest), Co, Cu, and Pb (only for Vineyard vs Forest). Elements such as Al, As, Co and Cu show large positive log-differences with Vineyard concentrations exceeding those at Forest, Suburban and Urban sites. By contrast, Cd presents significantly higher concentrations at Urban relative to Vineyard, Suburban or Forest sites and Mn was markedly higher in Forest bees, showing notable site-specific variations.

**Figure 4.**
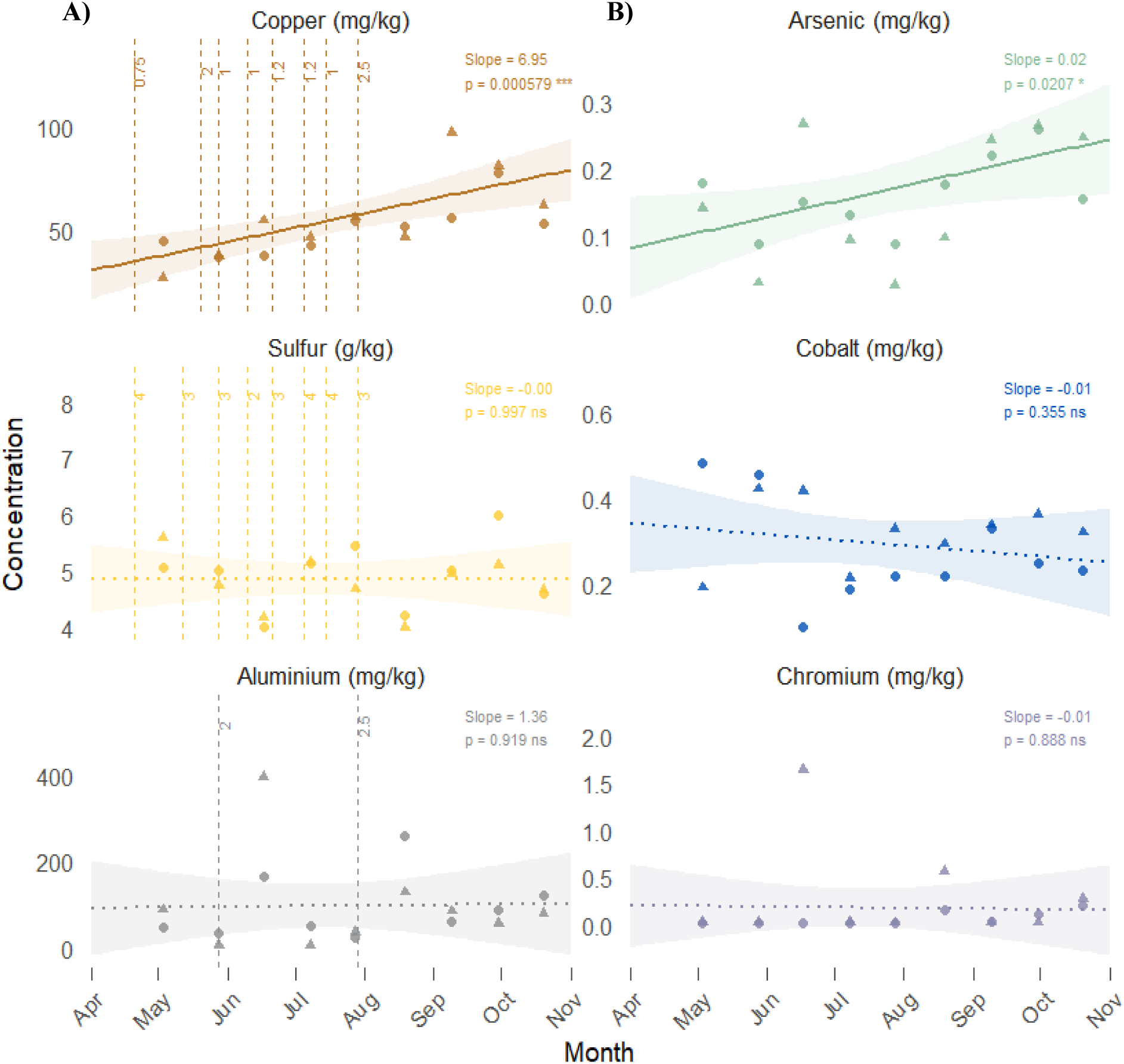
Monthly evolution of A) copper, sulfur and aluminum concentrations in honey bees and doses of phytosanitary treatments applied at the nearest vineyard plot and B) arsenic, cobalt and chromium concentrations in honey bees from the vineyard location. Individual points represent the two replicate hives. Solid lines represent significant temporal trends (p < 0.05) and dotted lines represent non-significant trends, as estimated by linear mixed-effect models with hive as a random intercept. Dashed vertical lines represent treatment doses in kg per ha.

Major elements such as Calcium (Ca), Phosphorous (P), Potassium (K), Magnesium (Mg), Sodium (Na), and Sulfur (S) did not show any significant differences between the study sites. They generally exhibited moderate levels across all sites, with notable peaks in the Vineyard location, especially for K and Mg during late summer, corresponding to periods of vineyard fertilization. Concentrations of potentially toxic metals such as Pb, Co, Cd or As were generally low and did not exceed a threshold of concern. Concentrations of elements at each sampling date and location can be found in Supporting Information (Supporting Information, Fig. S1).

The temporal evolution of the elements Cu, Al, As, Co and Cr found in higher concentrations in the Vineyard location are displayed in Figure 4. Each dot represents measured concentrations at individual sampling dates, while fitted lines show the overall monthly trends. Graphics on the left (Fig. 4A) show the concentrations of Cu, S and Al in honey bees, three compounds recorded as phytosanitary treatments applied to the nearest vineyard plot. Only honey bees from the Vineyard site exhibited a significant increase in copper concentration over time (Fig. 4 and Supporting Information Fig. S2), with a pronounced upward slope (estimate: 7 ± 2 mg.kg^-1^ per month, p < 0.001) in concordance with repeated treatments (8 across the season). Concentrations of Cu continue to increase in the sampled honey bees even after the latest treatment applied on late July. In contrast, copper concentrations in bees sampled from Forest, Suburban, and Urban locations remained relatively stable, showing no temporal accumulation as indicated by non-significant slope estimates (Supporting Information, Fig. S2).

Sulfur concentrations remain relatively stable with minor fluctuations despite treatments applied on the vineyard plot (8 across the season). On the contrary, Al concentrations show higher levels than other sites (Supporting Information, Fig. S2) and two peaks in sampled honey bees after each of the two treatments against downy mildew (fosetyl-aluminium) applied on the nearest plot (lower panel Fig. 4A). This result highlights that high levels of Al can be reached in honey bees located close to vineyards after treatments with aluminium-containing pesticides. Levels of As, Co and Cr elements remain low (below 0.3, 0.5 and 1 mg kg⁻¹ respectively) even in the Vineyard location, the most contaminated study site. A significant upward temporal trend in As concentrations was detected over the sampling period (p < 0.05), suggesting a progressive accumulation in bees. In contrast, no significant temporal trends were identified for Co and Cr concentrations although Cr concentrations in honey bees show a temporal co-occurrence and similar profile than Al.

### Pesticide residues

Figure 5 shows the temporal and across-site distribution of pesticide residues detected in the honey bees sampled across four different land use types: Forest, Suburban, Urban, and Vineyard. Out of the 57 molecules monitored with our analytical method (31 fungicides, 18 insecticides and acaricides, 7 herbicides, and one synergist) 14 fungicides and 2 acaricides were detected in our bee samples.

**Figure 5.**
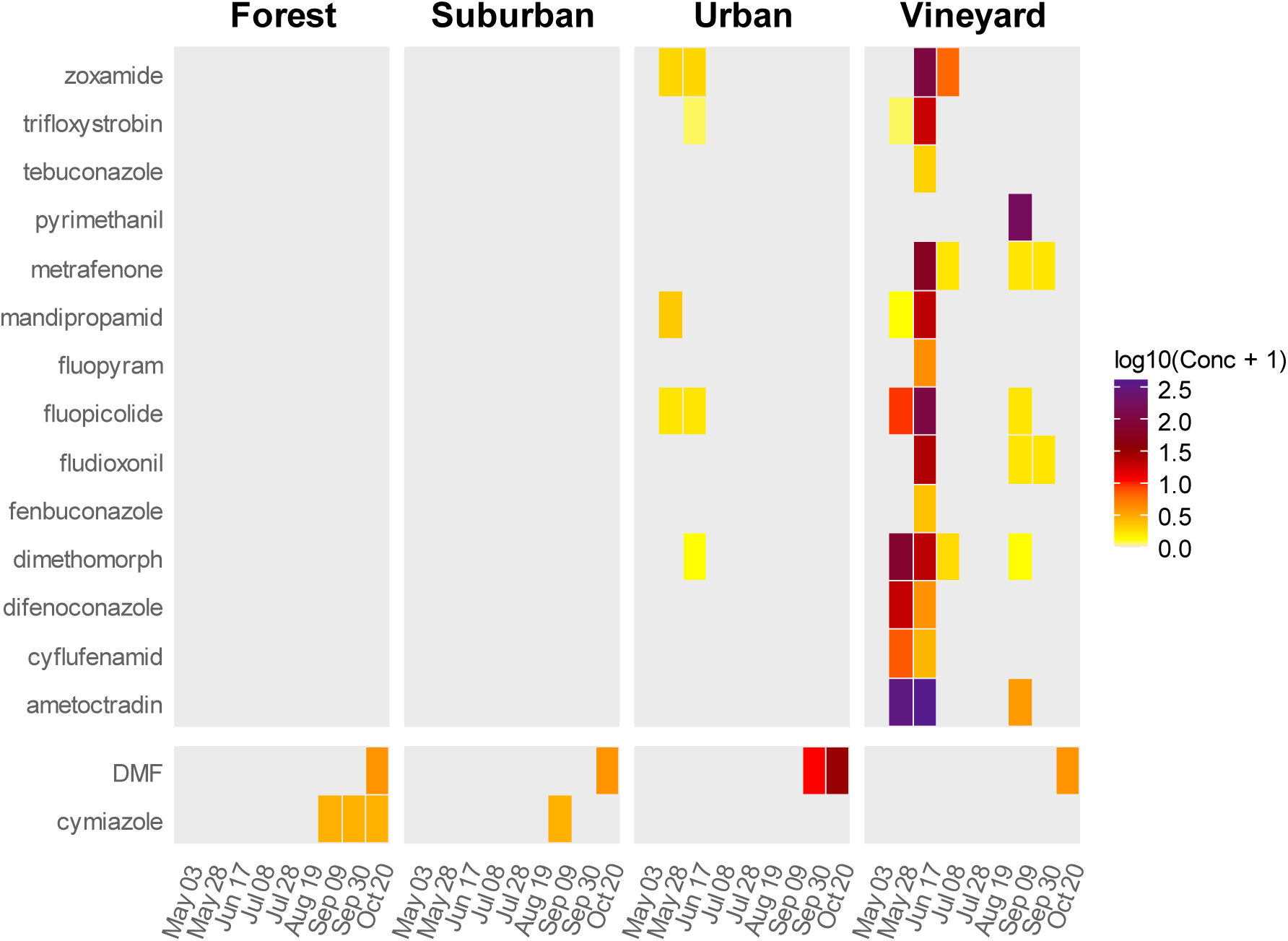
Heatmap of pesticide residues concentrations (ng.g^-1^) quantified in honey bees sampled between May and October in contrasted locations (Forest, Suburban, Urban, Vineyard). Data presented are log₁₀-transformed maximum concentrations (between the two sampled hives), fungicides on the upper part (14) and acaricides (2) on the lower part of the map. Color intensity indicating relative abundance (from yellow = low to purple = high concentrations).

Fungicide richness varied significantly across locations (Friedman test: χ² = 13.5, df = 3, p < 0.01). Post-hoc pairwise comparisons revealed that the Vineyard location harboured significantly higher fungicide richness than both Forest (p < 0.01) and Suburban (p < 0.01) locations, while no significant difference was detected between Vineyard and Urban locations (p > 0.05) nor between Forest, Suburban and Urban locations (p > 0.05). No significant temporal trend in fungicide richness was detected in any location over the sampling period (Spearman correlation, all p > 0.05). The Urban location demonstrated a broad spectrum of fungicide residues, with multiple compounds including zoxamide, trifloxystrobin, mandipropamid, fluopicolide and dimetomorph detected intermittently from May to September at low levels across a few sampling dates. This is expected, since vineyard plots are also present in the vicinities of the sampled hives at INRAE research center (Fig.1), explaining why common fungicides are also found in honey bees. In contrast, bees from the Vineyard site exhibited the highest number, frequency and concentration of fungicide residues with 14 different fungicides detected across multiple sampling dates at concentrations sometimes exceeding log₁₀(Conc + 1) of 2.5 (Fig. 5). This corroborates intensive fungicide use in vineyards, particularly between May and July. Two acaricides were found: DMF (2,4-diméthylformamidine) and cymiazole. DMF is a metabolite of amitraz, an acaricide that was used to treat our honey bee colonies against *Varroa destructor*, usually applied in hives at the end of September, justifying the observed peak of concentration. Cymiazole was present in bees from the Forest and Suburban locations however it was not used in our colonies to treat against Varroa mites. This substance is frequently used as an antiparasitic in veterinary medicine against external parasites (lice, mites, ticks) of livestock and domestic animals.

As for elements, correspondence between quantified pesticide residues in bees from the Vineyard location and phytosanitary products used to treat the nearest vineyard plot was investigated (Fig. 6). This figure shows the concentrations of the 14 pesticide residues measured in bees at the sampling dates and visualization of how treatments correspond to residues following field applications. It was observed that the bees were also contaminated by certain active substances that were not listed in the treatment plan provided by the wine grower (bars filled with diagonal lines). These substances are fungicides used to control cryptogamic diseases such as powdery mildew, downy mildew, and *Botrytis cinerea* and have very probably been used in other vineyard plots close to the sampled hives. Out of the 14 different fungicides quantified in the sampled honey bees, 8 can be linked to a field application on the nearest vineyard plot. These results demonstrate contamination events in bees following pesticide applications, which might lead to sub-lethal or chronic effects at the individual and colony levels. The highest concentrations appeared in June, right after several treatments, with multiple pesticides residues detected simultaneously and a cumulative concentration of pesticide residues reaching more than 800 ng.g**^- 1^**. Different pesticides contributed variably: some (e.g., ametoctradin, fluopicolide, dimetomorph, pyrimethanil, zoxamide) dominated the residue profile at specific dates and ametoctradin was shown to contribute to more than half of the total amount of residues quantified in May and June. Together with fluopicolide, ametoctradin was quantified in higher proportions on the second sampling (June 17th) than on the first sampling (May 28th) after application in the field. In one case, an active substance (metrafenone, June 17th) was quantified in honey bees prior to the date of treatment given by the wine grower. This could be explained if another plot nearby was treated with the substance prior to sampling or if the wine grower made a mistake in the calendar.

**Figure 6.**
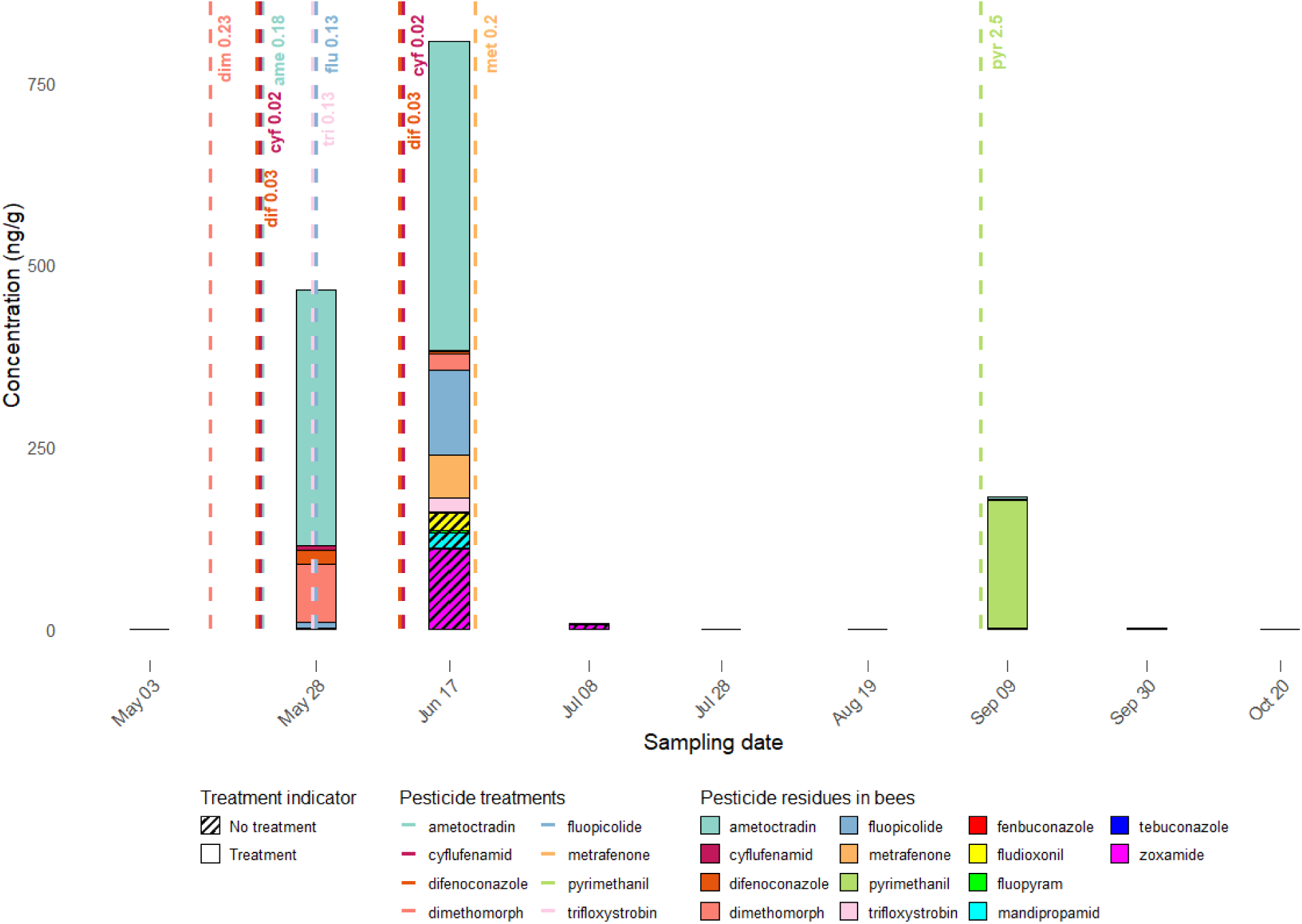
Seasonal changes in pesticide residues concentrations in honey bees from the vineyard location and associated pesticide applications in the vineyard. Bars represent maximum concentrations (between the two sampled hives) quantified in ng.g^-1^ and dashed lines indicate the dates of treatments and the dose in g of active substance per ha applied on the nearest plot obtained from the wine grower. Striped colors represent pesticide residues found in honey bees but not applied on the nearest plot.

To explore potential co-exposure to copper and synthetic fungicides in vineyard bees, we examined the co-occurrence of Cu and synthetic fungicide residues throughout the season (Figure S3, Spearman, *ns*). Notably, the highest Cu concentrations (ca. 80–100 mg/kg) co-occurred with detections of pyrimethanil, fludioxonil, metrafenone, dimethomorph and fluopicolide, while ametoctradin was detected at high concentrations (> 400 ng/g) at intermediate Cu levels (ca. 35–55 mg/kg). Across all sampling dates, at least one fungicide was co-detected with Cu in the majority of vineyard bee samples (Figure S3).

## Discussion

### Honey bees as biosentinels of different environmental contexts

This study shows that honey bee colonies settled in contrasted environments present different contamination profiles that can be related to specific environments and practices, showing their reliability as biosentinels of environmental contaminants. We show that contamination by synthetic pesticides residues and elements peaks in the agriculturally managed habitat, which is the Vineyard environment. The temporal pattern of contamination indicates higher concentrations during the growing season (May–July), coinciding with pesticide application periods in vineyards and leaving a distinct signature on honey bees under natural conditions.

Historical and current agricultural practices, particularly the widespread use of copper-based fungicides and arsenic-containing agrochemicals since the late 19th century, have led to a significant accumulation of these elements in vineyard soils which can then enter the food chains via plants (Bertoldi et al., 2013; Scott & Gardiner, 2025). Honey bees and other wild bees are highly present and active in vineyards (Kratschmer et al., 2019) particularly attracted by ground vegetation cover and flower diversity (Biella et al., 2025). However, this exposes them to pesticide residues and elements collected from floral resources (Botías et al., 2015; McArt et al., 2017; Xun et al., 2017, 2018; Mattiello et al., 2023; Scott & Gardiner, 2025) as well as to direct exposure during spraying. Levels of several elements measured in honey bees in the Vineyard location exceed the levels found in other studies (van der Steen et al., 2012). Concentrations of Cu have been reported in honey bee samples from 5 mg kg^−1^ to around 20 mg kg^−1^ in urban areas (Salvaggio et al., 2017; Giglio et al., 2017; Skorbiłowicz et al., 2018). In this study, Cu levels in honey bees reached a maximum of 97 mg kg^−1^ in the Vineyard location and showed temporal accumulation. Elements such as Cu and As both tend to accumulate in the surface horizons of soils because their natural depletion and removal occurs only very slowly (Aide et al., 2016; Sereni et al., 2024). Arsenic concentrations measured in honey bees were higher in the Vineyard location and showed a significant temporal accumulation in honey bees but did not reach a threshold of concern in comparison to the levels found in other studies (Giglio et al., 2017; Salvaggio et al., 2017). Just like Cu, Mn is an essential element that supports many processes in plants and organisms but its presence in excess can be deleterious (Søvik et al., 2015). It is to be noted that we found Mn in significantly higher proportions in honey bees from the Forest location. In the surface layers of forests, the more acidic soils have the higher concentrations of Mn originating from the standing leaves and litterfall of trees (Zaitsev et al., 2020; Michopoulos et al., 2021). Heathers (genus Calluna and Erica) were the main flower resources of the two colonies in the Forest location and the capacity of these plants to accumulate this element (Kula et al., 2012) could be the source of Mn found in bees. In urban areas, higher levels of Cd were quantified in honey bees in comparison to the other locations due to historical industrial activity, traffic, and waste, but stayed within the range of concentrations found in other studies (Hernández-Medina et al., 2025). As soil samples were not collected in this study, this inevitably limits our ability to connect bee contamination levels with local soil concentrations or to separate past contamination from more recent inputs. Future studies should therefore include soil and other environmental samples to help clarify where these contaminants come from and how they move through the ecosystem.

While a great amount of attention has focused on certain insecticides, fungicides are typically the most abundant pesticide residues found in bumble bees (Botías et al., 2017), honey bees and hive material (Mullin et al., 2010; Sanchez-Bayo & Goka, 2014; Cappellari et al., 2024). Similar pattern occur in other species such as spiders or hornets, particularly when crops such as orchard or vineyards are nearby (Tison et al., 2023; Cappellari et al., 2024; Michalko et al., 2024). In this study, 14 different synthetic fungicides were detected in honey bees, with up to 13 detected simultaneously in June (Fig. 5 and 6), several of which are similar to those reported in the above-cited studies. Some fungicides dominated the residue profile such as ametoctradin, contributing to more than half of the total amount of residues quantified in May and June (Fig. 5). Ametoctradin, but also fluopicolide, the second most found substance, are being known to have a high probability of bioconcentration within organisms (Lewis et al., 2016). However, measured concentrations of synthetic fungicides in honey bees from the Vineyard location do not appear to exceed thresholds of concern based on known toxicological endpoints for honey bees, with LD50 values and regulatory thresholds reported in Supporting Information (Table S4).

Aluminium seems to peak in honey bees at high concentrations (up to 399 mg kg^-1^) in the Vineyard location after field applications before getting back to lower levels, however still higher than in other locations. Fosetyl-aluminium is a systemic fungicide currently used against downy mildew which has a high potential for contaminating pollinators resources but a low bioconcentration factor in organisms (Lewis et al., 2016). Cr concentrations in honey bees show a similar profile than Al, possibly suggesting mutual presence in fosetyl-aluminium fungicide treatment or temporal co-occurrence during fungicide spraying and associated vineyard operations increasing bee contact with Al-rich, Cr-bearing mineral material (soil dust, spray drift, contaminated water/foliage).

Sulfur concentrations remain relatively stable with minor fluctuations despite repeated treatments, suggesting that these applications only weakly affect the large, biologically regulated sulfur pool in bees. This reflects our measurement of total S, which integrates endogenous and dietary sulfur with any exogenous residues, and the fact that vineyard sulfur formulations are contact products with short persistence and high volatility.

On the contrary to Al, we found that Cu accumulates in honey bees along the viticultural season, even after the end of field treatment applications. A progressive Cu buildup in bees repeatedly exposed to Bordeaux mixture in vineyards was shown. However, without data on bee turnover or tissue distribution, true biological accumulation cannot be inferred. Chronic Cu contamination in honey bees likely reflects repeated re-exposure through surface adsorption and/or ingestion of environmental residues after successive treatments. It may also result from accumulation within colonies when young bees consume Cu-contaminated food. To our knowledge, this is the first evidence of temporal copper accumulation in non-target terrestrial organisms under natural conditions. Previous studies reported that average levels of Cu and trace elements were higher in bees than in nectar or honey, indicating that these elements can transfer in floral resources and bioaccumulate in bees (Hladun et al., 2015; Xun et al., 2017, 2018; Scott et al., 2024; Scott & Gardiner, 2025). These findings suggest that bees are better sentinels of environmental contamination than honey (Silici et al., 2016; Grzegorz et al., 2021).

### Exposure and risk assessment

Real-world exposure to fungicides alone or in combination with other substances and their impact on bees is still the subject of great uncertainty (Rondeau & Raine, 2022) as ecotoxicological studies involving fungicides and especially inorganic fungicides remain scarce. Bees are usually unable to detect or avoid pesticides residues or elements in their food (Monchanin et al., 2022; Parkinson et al., 2023; Gekière, Breuer, et al., 2024) which consequently exposes foragers and other colony members to these contaminants.

Copper exhibit higher maximum doses of application (about 20 times higher than for modern synthetic fungicides) and significantly higher risks for honey bees (3- to 20-fold) compared with modern synthetic fungicides used for the same purpose (Burandt et al., 2024). However, data on chronic toxicity endpoints and sublethal effects of Cu remain limited (European Food Safety Authority (EFSA) et al., 2018). A list of ecotoxicological endpoints and lethal dose 50 % (LD50) is available for some elements in Gekière et al. (2023, supplementary material). However, comparison with these ecotoxicological thresholds should be interpreted with caution, as our data do not differentiate between external and internal contamination given that bees were not washed prior to chemical analysis. The concentration of Cu quantified in honey bees from the Vineyard location exceeded several times the lethal concentration (LC50: concentration of Cu that killed 50 % of the tested honey bees) of 72 mg L^−1^ for foragers and 7 mg L^−1^ for larvae (Di et al., 2016). Altered feeding behavior and survival in adult honey bees fed with Cu were observed (Di et al., 2016; Burden et al., 2019). In other studies, the lethal dose (LD50) was 12 mg per bee for copper oxychloride and 49 mg per bee for copper hydroxide considering the ecotoxicological risk as moderate for bees (Lewis et al., 2016; European Food Safety Authority (EFSA) et al., 2018). Nonetheless, at the concentrations found in this study, one can expect substantial reductions in larval growth and survival (Di et al., 2016). If accumulation occurs following feeding, as suggested by our results, the impact could be severe and severe reduction in whole colony health could be expected.

Since bees screened for pesticides residues are collected alive, residues in bees most likely represent an underestimation of their true exposure. This is because bees that have been exposed to lethal doses are unlikely to be caught alive (Botías et al., 2017; Martinello et al., 2020). Although acute toxicity thresholds are usually not exceeded in this study except for Cu and Al, several organic fungicides pose chronic risks, especially for wild bees (Rondeau & Raine, 2022). Fungicides when applied correctly are usually not responsible for the direct death of bees but can have sublethal behavioral or physiological effects for all castes and at the colony level which could lead to a reduction in foraging activity or brood care (Bryden et al., 2013; Godfray et al., 2014; Berenbaum & Liao, 2019). Organic fungicides have also been shown to kill honey bee midgut cells and reduce beneficial fungi, increasing their susceptibility to pathogens and weakening colonies (Pettis et al., 2013; Yoder et al., 2017) whereas an exposure to Cu perturbated in the redox balance of bumble bees (Gekière, Gérard, et al., 2024). As for other elements, a single exposure to Al at concentrations as low as 2 and 20 mg L^-1^ negatively impacted honey bees’ floral choices (Chicas-Mosier et al., 2017) and exposure to As, Pb, Cu or combinations of these elements at concentrations far below the levels found in our study were shown to slow down appetitive learning and reduce long-term memory specificity in honey bees (Monchanin et al., 2021). Given that learning and memory of olfactory cues play crucial roles in the behavioral ecology of honey bees, acute or chronic exposure to elements could have important consequences on pollination (Klein et al., 2017). Furthermore, frequent fungicide–insecticide mixtures can increase the potential for harmful synergistic effects and should be taken seriously (Sanchez-Bayo & Goka, 2014; Traynor et al., 2016). This is also the case for supposedly inert ingredients in fungicide mixtures that can mediate acute effects on honey bee learning performance (DesJardins et al., 2023). Several fungicides, including pyrimethanil, ametoctradin, dimethomorph, and fluopicolide, were detected at substantial concentrations (up to > 400 ng/g) in the same bee samples as elevated Cu levels, suggesting that bees foraging in vineyard environments are routinely exposed to complex mixtures rather than individual compounds. This is consistent with viticultural practices in the study area, where Cu-based treatments are commonly applied alongside synthetic fungicides across the growing season. While our dataset does not allow us to formally test for synergistic or additive effects that could enhance toxicity, these observations support the concern that field-realistic exposure of vineyard bees to Cu or other elements cannot be considered in isolation from simultaneous fungicide contamination. For example, Cu itself exerts toxicity largely through oxidative stress. Several of the fungicides identified here such as azole fungicides (e.g., difenoconazole, fenbuconazole, tebuconazole) and some quinone outside inhibitors (QoI) (e.g., fluopicolide, ametoctradin) also modulate mitochondrial electron transport or cellular redox balance, which may amplify oxidative stress when combined with Cu exposure.

Our findings are consistent with the One Health framework and directly support recent policy priorities outlined in the EU Pollinators Initiative (European Commission, 2023), which emphasize the need to address cumulative stressors affecting pollinator health. By demonstrating the co-occurrence and potential combined effects of pesticides and trace elements, this study underscores the importance of moving beyond single-compound risk assessments toward more integrative approaches that better reflect real-world exposure conditions and inform effective mitigation strategies

### Perspectives

Pollinators and more generally arthropods represent an entry point for the transfer of contaminants in terrestrial food webs and could have cascading consequences on ecosystems (Pesce et al., 2024; Tison et al., 2024; Faburé et al., 2024). Several studies have shown the presence of contaminants including fungicides residues and elements in predators such as hornets, spiders and birds (Moreau et al., 2022; Tison et al., 2023; Michalko et al., 2024; Panda et al., 2025) with possible lethal or sublethal effects likely to propagate in food webs via trophic interactions. Positive Cu–Zn correlations across soil, prey, and feather samples illustrate trace metal transfer through trophic levels, affecting higher organisms like birds (Panda et al., 2025). Critical gaps in knowledge still exist, such as the effect of fungicides on secondary consumers (Faburé et al., 2024) or the impact of low levels of elements on arthropods and their predator cognitive abilities.

Pooling 10 foragers per sample, required to achieve sufficient biomass for pesticide residues and elements analysis, masks within-colony variability that could reveal exposure heterogeneity among individuals. Future studies using individual-level analyses or several pools of samples per colony would provide complementary insights into intra-colony exposure differences. Current risk assessment approaches however typically focus on the effects of individual elements or pesticides residues, overlooking the potential for interactions between substances (Gekière et al., 2023). This limitation is particularly relevant for honey bees and other species that share and store contaminated food, which can amplify these interactions within populations. Assessing the interactive effects of major and trace elements with pesticides residues (active ingredients, co-formulants and adjuvants) together with other environmental factors calls for further investigations (Singh et al., 2017; Sgolastra et al., 2018; Committee et al., 2019; Naqash et al., 2020; Gekière et al., 2023).

Another limitation of this study is that only parent pesticides residues were tracked and quantified, whereas they can be transformed in the environment or metabolized in bees. Therefore, pesticide degradation products should be monitored in future studies. Furthermore, because each land-cover category was represented by a single site, the observed differences in chemical profiles may also reflect site-specific conditions. Future studies including more sites within each land-cover type, and a broader range of land-cover compositions, would help strengthen conclusions about the relationships between land cover and chemical profiles in honey bees.

## Conclusion

Our study highlights the major impacts of land use and seasonal dynamics on contaminants distribution in honey bees. Consistent with our first hypothesis, environments with intensive chemical use, such as vineyards, leave a stronger ecological footprint on honey bees compared to habitats with low chemical input. This is reflected in the higher and more diverse contaminant loads observed at the vineyard site. In line with our second hypothesis, the Vineyard location stands out for several key elements and pesticide residues that reflect a distinct footprint. This footprint arises from combined exposure to both historical and current inputs of organic and inorganic compounds, including several fungicides whose effects are still understudied in insects. These results reveal distinct temporal and across-site patterns of pesticide residues and element concentrations in bees, reflecting localized anthropogenic influences and underscoring the utility of honey bees as environmental sentinels. The clear correlation between field applications and bee residue loads strengthens the necessity for monitoring and regulating application timing to protect pollinators. Finally, supporting our third hypothesis, copper shows significant accumulation trends in bees collected in vineyards, demonstrating for the first time, the temporal accumulation of copper in naturally exposed pollinators.

Given the tendency of elements and pesticides residues to transfer and accumulate in environmental matrices and organisms, it is absolutely necessary to consider contaminant bioaccumulation and their interactions in ecotoxicological assessments and future research (Gekière et al., 2023; Tison et al., 2024).

## Acknowledgements

We would like to thank the three reviewers from PCI Ecotoxicology and Environmental Chemistry for their valuable comments which greatly helped to improve the manuscript. We also thank the landowners at the Forest, Vineyard and Suburban locations for letting us install the beehives and access their land for sampling. We greatly valued Pierre Masson’s early contributions to this project and deeply regret his untimely passing.

## Data, scripts, code, and supplementary information availability

Data, script and data dictionary are available online: Tison, L. (2025). Data tables and R script [Data set]. Zenodo. https://doi.org/10.5281/zenodo.17951865. Supporting Information is available online: Tison, L. (2025). Supporting Information. Zenodo. https://doi.org/10.5281/zenodo.17940256. Raw concentrations of pesticides and elements can be visualized.

## References

Aide M, Beighley D, Dunn D (2016) Arsenic In The Soil Environment: A Soil Chemistry Review.

Albaseer SS, Jaspers VLB, Orsini L, Vlahos P, Al-Hazmi HE, Hollert H (2025) Beyond the field: How pesticide drift endangers biodiversity. Environmental Pollution, 366, 125526. 10.1016/j.envpol.2024.125526

Badiou-Bénéteau A, Benneveau A, Géret F, Delatte H, Becker N, Brunet JL, Reynaud B, Belzunces LP (2013) Honeybee biomarkers as promising tools to monitor environmental quality. Environment International, 60, 31–41. 10.1016/j.envint.2013.07.002

Bates D, Mächler M, Bolker B, Walker S (2015) Fitting Linear Mixed-Effects Models Using lme4. Journal of Statistical Software, 67, 1–48. 10.18637/jss.v067.i01

Beketov MA, Kefford BJ, Schäfer RB, Liess M (2013) Pesticides reduce regional biodiversity of stream invertebrates. Proceedings of the National Academy of Sciences, 110, 11039–11043. 10.1073/pnas.1305618110

Berenbaum MR, Liao L-H (2019) Honey Bees and Environmental Stress: Toxicologic Pathology of a Superorganism. Toxicologic Pathology, 47, 1076–1081. 10.1177/0192623319877154

Bertoldi D, Villegas TR, Larcher R, Santato A, Nicolini G (2013) Arsenic present in the soil-vine-wine chain in vineyards situated in an old mining area in Trentino, Italy. Environmental Toxicology and Chemistry, 32, 773–779. 10.1002/etc.2119

Bianco A, Nibert P, Wu Y, Baray J-L, Brigante M, Mailhot G, Deguillaume L, Vione D, Cabanes DJE, Méjean M, Besse-Hoggan P (2025) Are Clouds a Neglected Reservoir of Pesticides? Environmental Science & Technology, 59, 21579–21588. 10.1021/acs.est.5c03787

Biella P, Ramazzotti F, Parolo G, Galimberti A, Labra M, Brambilla M (2025) Vineyard footprint on pollinators is mediated by flower vegetation, organic farming, seasonal and weather factors, a case study from North Italy. Agriculture, Ecosystems & Environment, 378, 109297. 10.1016/j.agee.2024.109297

Botías C, David A, Hill EM, Goulson D (2017) Quantifying exposure of wild bumblebees to mixtures of agrochemicals in agricultural and urban landscapes. Environmental Pollution, 222, 73–82. 10.1016/j.envpol.2017.01.001

Botías C, David A, Horwood J, Abdul-Sada A, Nicholls E, Hill E, Goulson D (2015) Neonicotinoid Residues in Wildflowers, a Potential Route of Chronic Exposure for Bees. Environmental Science & Technology, 49, 12731–12740. 10.1021/acs.est.5b03459

Bryden J, Gill RJ, Mitton RAA, Raine NE, Jansen VAA (2013) Chronic sublethal stress causes bee colony failure. Ecology Letters, 16, 1463–1469. 10.1111/ele.12188

Burandt QC, Deising HB, von Tiedemann A (2024) Further Limitations of Synthetic Fungicide Use and Expansion of Organic Agriculture in Europe Will Increase the Environmental and Health Risks of Chemical Crop Protection Caused by Copper-Containing Fungicides. Environmental Toxicology and Chemistry, 43, 19–30. 10.1002/etc.5766

Burden CM, Elmore C, Hladun KR, Trumble JT, Smith BH (2016) Acute exposure to selenium disrupts associative conditioning and long-term memory recall in honey bees (*Apis mellifera*). Ecotoxicology and Environmental Safety, 127, 71–79. 10.1016/j.ecoenv.2015.12.034

Burden CM, Morgan MO, Hladun KR, Amdam GV, Trumble JJ, Smith BH (2019) Acute sublethal exposure to toxic heavy metals alters honey bee (Apis mellifera) feeding behavior. Scientific Reports, 9, 4253. 10.1038/s41598-019-40396-x

Calatayud-Vernich P, Calatayud F, Simó E, Picó Y (2018) Pesticide residues in honey bees, pollen and beeswax: Assessing beehive exposure. Environmental Pollution, 241, 106–114. 10.1016/j.envpol.2018.05.062

Cappellari A, Malagnini V, Fontana P, Zanotelli L, Tonidandel L, Angeli G, Ioriatti C, Marini L (2024) Impact of landscape composition on honey bee pollen contamination by pesticides: A multi-residue analysis. Chemosphere, 349, 140829. 10.1016/j.chemosphere.2023.140829

Chicas-Mosier AM, Cooper BA, Melendez AM, Pérez M, Oskay D, Abramson CI (2017) The effects of ingested aqueous aluminum on floral fidelity and foraging strategy in honey bees (*Apis mellifera)*. Ecotoxicology and Environmental Safety, 143, 80–86. 10.1016/j.ecoenv.2017.05.008

Committee ES, More SJ, Bampidis V, Benford D, Bennekou SH, Bragard C, Halldorsson TI, Hernández-Jerez AF, Koutsoumanis K, Naegeli H, Schlatter JR, Silano V, Nielsen SS, Schrenk D, Turck D, Younes M, Benfenati E, Castle L, Cedergreen N, Hardy A, Laskowski R, Leblanc JC, Kortenkamp A, Ragas A, Posthuma L, Svendsen C, Solecki R, Testai E, Dujardin B, Kass GE, Manini P, Jeddi MZ, Dorne J-LC, Hogstrand C (2019) Guidance on harmonised methodologies for human health, animal health and ecological risk assessment of combined exposure to multiple chemicals. EFSA Journal, 17, e05634. 10.2903/j.efsa.2019.5634

Cozmuta AM, Bretan L, Cozmuta LM, Nicula C, Peter A (2012) Lead traceability along soil-melliferous flora-bee family-apiary products chain. Journal of Environmental Monitoring, 14, 1622–1630. 10.1039/C2EM30084B

Cunningham SA, Crane MJ, Evans MJ, Hingee KL, Lindenmayer DB (2022) Density of invasive western honey bee (Apis mellifera) colonies in fragmented woodlands indicates potential for large impacts on native species. Scientific Reports, 12, 3603. 10.1038/s41598-022-07635-0

Cunningham MM, Tran L, McKee CG, Ortega Polo R, Newman T, Lansing L, Griffiths JS, Bilodeau GJ, Rott M, Marta Guarna M (2022) Honey bees as biomonitors of environmental contaminants, pathogens, and climate change. Ecological Indicators, 134, 108457. 10.1016/j.ecolind.2021.108457

Decourtye A, Rollin O, Requier F, Allier F, Rüger C, Vidau C, Henry M (2023) Decision-making criteria for pesticide spraying considering the bees’ presence on crops to reduce their exposure risk. Frontiers in Ecology and Evolution, 11, 1062441. 10.3389/fevo.2023.1062441

DesJardins NS, Harrison JF, Smith BH (2023) The effects of anthropogenic toxins on honey bee learning: Research trends and significance. Apidologie, 54, 59. 10.1007/s13592-023-01040-w

Di N, Hladun KR, Zhang K, Liu T-X, Trumble JT (2016) Laboratory bioassays on the impact of cadmium, copper and lead on the development and survival of honeybee (Apis mellifera L.) larvae and foragers. Chemosphere, 152, 530–538. 10.1016/j.chemosphere.2016.03.033

Di Prisco G, Cavaliere V, Annoscia D, Varricchio P, Caprio E, Nazzi F, Gargiulo G, Pennacchio F (2013) Neonicotinoid clothianidin adversely affects insect immunity and promotes replication of a viral pathogen in honey bees. Proceedings of the National Academy of Sciences of the United States of America, 110, 18466–18471. 10.1073/pnas.1314923110

European Commission. (2023). Revision of the EU Pollinators Initiative – A new deal for pollinators. COM (2023) 35 final, 24 January 2023. Available at: https://eur-lex.europa.eu/legal-content/EN/ALL/?uri=COM:2023:35:FIN

EFSA Scientific Committee, More S, Bampidis V, Benford D, Bragard C, Halldorsson T, Hernández-Jerez A, Bennekou SH, Koutsoumanis K, Machera K, Naegeli H, Nielsen SS, Schlatter J, Schrenk D, Silano V, Turck D, Younes M, Arnold G, Dorne J, Maggiore A, Pagani S, Szentes C, Terry S, Tosi S, Vrbos D, Zamariola G, Rortais A (2021) A systems-based approach to the environmental risk assessment of multiple stressors in honey bees. EFSA Journal, 19. 10.2903/j.efsa.2021.6607

European Food Safety Authority (EFSA), Arena M, Auteri D, Barmaz S, Bellisai G, Brancato A, Brocca D, Bura L, Byers H, Chiusolo A, Court Marques D, Crivellente F, De Lentdecker C, Egsmose M, Erdos Z, Fait G, Ferreira L, Goumenou M, Greco L, Ippolito A, Istace F, Jarrah S, Kardassi D, Leuschner R, Lythgo C, Magrans JO, Medina P, Miron I, Molnar T, Nougadere A, Padovani L, Parra Morte JM, Pedersen R, Reich H, Sacchi A, Santos M, Serafimova R, Sharp R, Stanek A, Streissl F, Sturma J, Szentes C, Tarazona J, Terron A, Theobald A, Vagenende B, Verani A, Villamar-Bouza L (2018) Peer review of the pesticide risk assessment of the active substance copper compounds copper(I), copper(II) variants namely copper hydroxide, copper oxychloride, tribasic copper sulfate, copper(I) oxide, Bordeaux mixture. EFSA Journal, 16. 10.2903/j.efsa.2018.5152

Faburé J, Hedde M, Le Perchec S, Pesce S, Sucré E, Fritsch C (2024) Role of trophic interactions in transfer and cascading impacts of plant protection products on biodiversity: a literature review. Environmental Science and Pollution Research. 10.1007/s11356-024-35190-w

Gekière A, Breuer L, Dorio L, Vanderplanck M, Michez D (2024) Lethal effects and sex-specific tolerance of copper and cadmium in the buff-tailed bumble bee. Environmental Toxicology and Pharmacology, 110, 104546. 10.1016/j.etap.2024.104546

Gekière A, Gérard M, Evrard D, Breuer L, Dorio L, Maesen P, Vanderplanck M, Michez D (2024) Copper-accelerated pupation in larvae of the buff-tailed bumble bee. Apidologie, 56, 4. 10.1007/s13592-024-01134-z

Gekière A, Vanderplanck M, Michez D (2023) Trace metals with heavy consequences on bees: A comprehensive review. Science of The Total Environment, 895, 165084. 10.1016/j.scitotenv.2023.165084

Giglio A, Ammendola A, Battistella S, Naccarato A, Pallavicini A, Simeon E, Tagarelli A, Giulianini PG (2017) Apis mellifera ligustica, Spinola 1806 as bioindicator for detecting environmental contamination: a preliminary study of heavy metal pollution in Trieste, Italy. Environmental Science and Pollution Research, 24, 659–665. 10.1007/s11356-016-7862-z

Girotti S, Ghini S, Ferri E, Bolelli L, Colombo R, Serra G, Porrini C, Sangiorgi S (2020) Bioindicators and biomonitoring: honeybees and hive products as pollution impact assessment tools for the Mediterranean area. Euro-Mediterranean Journal for Environmental Integration, 5, 62. 10.1007/s41207-020-00204-9

Glinski DA, Purucker ST, Minucci JM, Richardson RT, Lin C-H, Johnson RM, Henderson WM (2024) Analysis of contaminant residues in honey bee hive matrices. Science of The Total Environment, 954, 176329. 10.1016/j.scitotenv.2024.176329

Godfray HCJ, Blacquière T, Field LM, Hails RS, Petrokofsky G, Potts SG, Raine NE, Vanbergen AJ, McLean AR (2014) A restatement of the natural science evidence base concerning neonicotinoid insecticides and insect pollinators. Proceedings of the Royal Society B: Biological Sciences, 281, 20140558. 10.1098/rspb.2014.0558

Grzegorz B, Sulborska A, Stawiarz E, Olszewski K, Wiącek D, Ramzi N, Nawrocka A, Jedryczka M (2021) Capacity of honeybees to remove heavy metals from nectar and excrete the contaminants from their bodies. Apidologie, 52. 10.1007/s13592-021-00890-6

Gutiérrez M, Molero R, Gaju M, van der Steen J, Porrini C, Ruiz JA (2015) Assessment of heavy metal pollution in Córdoba (Spain) by biomonitoring foraging honeybee. Environmental Monitoring and Assessment, 187, 651. 10.1007/s10661-015-4877-8

Hernández-Medina ME, Montiel Pimentel JV, Castellanos I, Zuria I, Sánchez-Rojas G, Gaytán Oyarzun JC (2025) Metal concentration in honeybees along an urbanization gradient in Central Mexico. Environmental Research, 264, 120199. 10.1016/j.envres.2024.120199

Hladun KR, Di N, Liu T-X, Trumble JT (2015) Metal contaminant accumulation in the hive: Consequences for whole-colony health and brood production in the honey bee ( *Apis mellifera* L.). Environmental Toxicology and Chemistry, 35, 322–329. 10.1002/etc.3273

Klein S, Cabirol A, Devaud J-M, Barron AB, Lihoreau M (2017) Why Bees Are So Vulnerable to Environmental Stressors. Trends in Ecology & Evolution, 32, 268–278. 10.1016/j.tree.2016.12.009

Kratschmer S, Pachinger B, Schwantzer M, Paredes D, Guzmán G, Goméz JA, Entrenas JA, Guernion M, Burel F, Nicolai A, Fertil A, Popescu D, Macavei L, Hoble A, Bunea C, Kriechbaum M, Zaller JG, Winter S (2019) Response of wild bee diversity, abundance, and functional traits to vineyard inter-row management intensity and landscape diversity across Europe. Ecology and Evolution, 9, 4103–4115. 10.1002/ece3.5039

Kula E, Hrdlička P, Hedbávný J, Švec P (2012) Various content of manganese in selected forest tree species and plants in the undergrowth. Beskydy, 5, 19–26. 10.11118/beskyd201205010019

Kuznetsova A, Brockhoff PB, Christensen RHB (2017) lmerTest Package: Tests in Linear Mixed Effects Models. Journal of Statistical Software, 82, 1–26. 10.18637/jss.v082.i13

Lambert O, Piroux M, Puyo S, Thorin C, Larhantec M, Delbac F, Pouliquen H (2012) Bees, honey and pollen as sentinels for lead environmental contamination. Environmental Pollution, 170, 254–259. 10.1016/j.envpol.2012.07.012

Lewis KA, Tzilivakis J, Warner DJ, Green A (2016) An international database for pesticide risk assessments and management. Human and Ecological Risk Assessment: An International Journal, 22, 1050–1064. 10.1080/10807039.2015.1133242

Li N, Wang J, Song W-Y (2016) Arsenic Uptake and Translocation in Plants. Plant and Cell Physiology, 57, 4–13. 10.1093/pcp/pcv143

Lorenz S, Trau FN, Ruf LC, Meinikmann K, Fisch K, Stähler M, Schenke D, Blevins HL, Heinz M (2025) Pesticide contamination of small standing water bodies in the agricultural landscape of northeast Germany. Science of The Total Environment, 975, 179250. 10.1016/j.scitotenv.2025.179250

Mair KS, Irrgeher J, Haluza D (2023) Elucidating the Role of Honey Bees as Biomonitors in Environmental Health Research. Insects, 14, 874. 10.3390/insects14110874

Martinello M, Manzinello C, Borin A, Avram LE, Dainese N, Giuliato I, Gallina A, Mutinelli F (2020) A Survey from 2015 to 2019 to Investigate the Occurrence of Pesticide Residues in Dead Honeybees and Other Matrices Related to Honeybee Mortality Incidents in Italy. Diversity, 12, 15. 10.3390/d12010015

Mattiello A, Novello N, Cornu J-Y, Babst-Kostecka A, Pošćić F (2023) Copper accumulation in five weed species commonly found in the understory vegetation of Mediterranean vineyards. Environmental Pollution, 329, 121675. 10.1016/j.envpol.2023.121675

McArt SH, Urbanowicz C, McCoshum S, Irwin RE, Adler LS (2017) Landscape predictors of pathogen prevalence and range contractions in US bumblebees. Proceedings of the Royal Society B: Biological Sciences, 284, 20172181. 10.1098/rspb.2017.2181

Michalko R, Purchart L, Hofman J, Košulič O (2024) Distribution of pesticides in agroecosystem food webs differ among trophic groups and between annual and perennial crops. Agronomy for Sustainable Development, 44, 13. 10.1007/s13593-024-00950-y

Michopoulos P, Kostakis M, Thomaidis NS, Pasias I (2021) The influence of forest types on manganese content in soils. Folia Forestalia Polonica, 63, 1–9. 10.2478/ffp-2021-0001

Monchanin C, Drujont E, Devaud J-M, Lihoreau M, Barron AB (2021) Metal pollutants have additive negative effects on honey bee cognition. Journal of Experimental Biology, 224, jeb241869. 10.1242/jeb.241869

Monchanin C, Gabriela de Brito Sanchez M, Lecouvreur L, Boidard O, Méry G, Silvestre J, Le Roux G, Baqué D, Elger A, Barron AB, Lihoreau M, Devaud J-M (2022) Honey bees cannot sense harmful concentrations of metal pollutants in food. Chemosphere, 297, 134089. 10.1016/j.chemosphere.2022.134089

Moreau J, Rabdeau J, Badenhausser I, Giraudeau M, Sepp T, Crépin M, Gaffard A, Bretagnolle V, Monceau K (2022) Pesticide impacts on avian species with special reference to farmland birds: a review. Environmental Monitoring and Assessment, 194. 10.1007/s10661-022-10394-0

Mullin CA, Frazier M, Frazier JL, Ashcraft S, Simonds R, vanEngelsdorp D, Pettis JS (2010) High Levels of Miticides and Agrochemicals in North American Apiaries: Implications for Honey Bee Health. PLOS ONE, 5, e9754. 10.1371/journal.pone.0009754

Naqash N, Prakash S, Kapoor D, Singh R (2020) Interaction of freshwater microplastics with biota and heavy metals: a review. Environmental Chemistry Letters, 18, 1813–1824. 10.1007/s10311-020-01044-3

Ondrasek G, Shepherd J, Rathod S, Dharavath R, Imtiaz Rashid M, Brtnicky M, Shafiq Shahid M, Horvatinec J, Rengel Z (2025) Metal contamination – a global environmental issue: sources, implications & advances in mitigation. RSC Advances, 15, 3904–3927. 10.1039/D4RA04639K

Pamminger T, Botías C, Goulson D, Hughes WOH (2018) A mechanistic framework to explain the immunosuppressive effects of neurotoxic pesticides on bees. Functional Ecology, 32, 1921–1930. 10.1111/1365-2435.13119

Panda BP, Kumari R, Pradhan A, Majhi BK, Acharya P, Parida SP (2025) Assessing the bioaccumulation of copper and zinc using bird as bioindicator in different wetland ecosystems of Odisha, India. Environmental Systems Research, 14, 10. 10.1186/s40068-025-00404-8

Parkinson RH, Scott J, Dorling AL, Jones H, Haslam M, McDermott-Roberts AE, Wright GA (2023) Mouthparts of the bumblebee (Bombus terrestris) exhibit poor acuity for the detection of pesticides in nectar. eLife, 12. 10.7554/eLife.89129.2

Perugini M, Manera M, Grotta L, Abete MC, Tarasco R, Amorena M (2011) Heavy metal (Hg, Cr, Cd, and Pb) contamination in urban areas and wildlife reserves: honeybees as bioindicators. Biological Trace Element Research, 140, 170–176. 10.1007/s12011-010-8688-z

Pesce S, Mamy L, Sanchez W, Artigas J, Bérard A, Betoulle S, Chaumot A, Coutellec M-A, Crouzet O, Faburé J, Hedde M, Leboulanger C, Margoum C, Martin-Laurent F, Morin S, Mougin C, Munaron D, Nélieu S, Pelosi C, Leenhardt S (2024) The use of copper as plant protection product contributes to environmental contamination and resulting impacts on terrestrial and aquatic biodiversity and ecosystem functions. Environmental Science and Pollution Research. 10.1007/s11356-024-32145-z

Pettis JS, Lichtenberg EM, Andree M, Stitzinger J, Rose R, vanEngelsdorp D (2013) Crop Pollination Exposes Honey Bees to Pesticides Which Alters Their Susceptibility to the Gut Pathogen Nosema ceranae. PLOS ONE, 8, e70182. 10.1371/journal.pone.0070182

Porrini C, Caprio E, Tesoriero D, Prisco GD Using honey bee as bioindicator of chemicals in Campanian agroecosystems (South Italy)., 10.

R Core Team. (2025) R A Language and Environment for Statistical Computing. R Foundation for Statistical Computing. - References - Scientific Research Publishing

Raine NE, Gill RJ (2015) Tasteless pesticides affect bees in the field. Nature, 521, 38–39. 10.1038/nature14391

Rigal S, Dakos V, Alonso H, Auniņš A, Benkő Z, Brotons L, Chodkiewicz T, Chylarecki P, de Carli E, del Moral JC, Domşa C, Escandell V, Fontaine B, Foppen R, Gregory R, Harris S, Herrando S, Husby M, Ieronymidou C, Jiguet F, Kennedy J, Klvaňová A, Kmecl P, Kuczyński L, Kurlavičius P, Kålås JA, Lehikoinen A, Lindström Å, Lorrillière R, Moshøj C, Nellis R, Noble D, Eskildsen DP, Paquet J-Y, Pélissié M, Pladevall C, Portolou D, Reif J, Schmid H, Seaman B, Szabo ZD, Szép T, Florenzano GT, Teufelbauer N, Trautmann S, van Turnhout C, Vermouzek Z, Vikstrøm T, Voříšek P, Weiserbs A, Devictor V (2023) Farmland practices are driving bird population decline across Europe. Proceedings of the National Academy of Sciences, 120, e2216573120. 10.1073/pnas.2216573120

Rigal S, Perrot T (2025) Pesticides in France: ten years of combined exposure to active substances in land, air and surface water. Scientific Data, 12, 512. 10.1038/s41597-025-04864-6

Rondeau S, Raine NE (2022) Fungicides and bees: a review of exposure and risk. Environment International, 165, 107311. 10.1016/j.envint.2022.107311

Rueppell O, Bachelier C, Fondrk MK, Page RE (2007) Regulation of life history determines lifespan of worker honey bees (Apis mellifera L.). Experimental Gerontology, 42, 1020–1032. 10.1016/j.exger.2007.06.002

Ruschioni S, Riolo P, Minuz RL, Stefano M, Cannella M, Porrini C, Isidoro N (2013) Biomonitoring with Honeybees of Heavy Metals and Pesticides in Nature Reserves of the Marche Region (Italy). Biological Trace Element Research, 154, 226–233. 10.1007/s12011-013-9732-6

Salvaggio A, Pecoraro R, Scalisi EM, Tibullo D, Lombardo BM, Messina G, Loreto F, Copat C, Ferrante M, Avola R, D’amante G, Genovese C, Raccuia SA, Brundo MV (2017) Morphostructural and immunohistochemical study on the role of metallothionein in the detoxification of heavy metals in Apis mellifera L., 1758. Microscopy Research and Technique, 80, 1215–1220. 10.1002/jemt.22919

Sanchez-Bayo F, Goka K (2014) Pesticide Residues and Bees – A Risk Assessment (RNC Guedes, Ed,). PLoS ONE, 9, e94482. 10.1371/journal.pone.0094482

Scheringer M, Schulz R (2025) The State of the World’s Chemical Pollution.

Scott SB, Gardiner MM (2025) Trace Metals in Nectar of Important Urban Pollinator Forage Plants: A Direct Exposure Risk to Pollinators and Nectar-Feeding Animals in Cities. Ecology and Evolution, 15, e71238. 10.1002/ece3.71238

Scott SB, Lanno R, Gardiner MM (2024) Acute toxicity and bioaccumulation of common urban metals in Bombus impatiens life stages. Science of The Total Environment, 915, 169997. 10.1016/j.scitotenv.2024.169997

Sereni L, Paris J-M, Lamy I, Guenet B (2024) Estimations of soil metal accumulation or leaching potentials under climate change scenarios: the example of copper on a European scale. SOIL, 10, 367–380. 10.5194/soil-10-367-2024

Sgolastra F, Blasioli S, Renzi T, Tosi S, Medrzycki P, Molowny-Horas R, Porrini C, Braschi I (2018) Lethal effects of Cr(III) alone and in combination with propiconazole and clothianidin in honey bees. Chemosphere, 191, 365–372. 10.1016/j.chemosphere.2017.10.068

Silici S, Uluozlu OD, Tuzen M, Soylak M (2016) Honeybees and honey as monitors for heavy metal contamination near thermal power plants in Mugla, Turkey. Toxicology and Industrial Health, 32, 507–516. 10.1177/0748233713503393

Singh N, Gupta VK, Kumar A, Sharma B (2017) Synergistic Effects of Heavy Metals and Pesticides in Living Systems. Frontiers in Chemistry, 5, 70. 10.3389/fchem.2017.00070

Siviter H, Koricheva J, Brown MJF, Leadbeater E (2018) Quantifying the impact of pesticides on learning and memory in bees (M Pocock, Ed,). Journal of Applied Ecology, 55, 2812–2821. 10.1111/1365-2664.13193

Skorbiłowicz E, Skorbiłowicz M, Cieśluk I (2018) Bees as bioindicators of environmental pollution with metals in an urban area. Journal of Ecological Engineering, 19, 229–234.

Søvik E, Perry CJ, LaMora A, Barron AB, Ben-Shahar Y (2015) Negative impact of manganese on honeybee foraging. Biology Letters, 11, 20140989. 10.1098/rsbl.2014.0989

Sponsler DB, Grozinger CM, Hitaj C, Rundlöf M, Botías C, Code A, Lonsdorf EV, Melathopoulos AP, Smith DJ, Suryanarayanan S, Thogmartin WE, Williams NM, Zhang M, Douglas MR (2019) Pesticides and pollinators: A socioecological synthesis. Science of The Total Environment, 662, 1012–1027. 10.1016/j.scitotenv.2019.01.016

Stanley J, Sah K, Jain SK, Bhatt JC, Sushil SN (2015) Evaluation of pesticide toxicity at their field recommended doses to honeybees, Apis cerana and A. mellifera through laboratory, semi-field and field studies. Chemosphere, 119, 668–674. 10.1016/j.chemosphere.2014.07.039

van der Steen JJM, de Kraker J, Grotenhuis T (2012) Spatial and temporal variation of metal concentrations in adult honeybees (Apis mellifera L.). Environmental Monitoring and Assessment, 184, 4119–4126. 10.1007/s10661-011-2248-7

Stemmelen A, Le Provost G, Giffard B, Rusch A (2025) Pesticide use and large patch size reduce natural pest control potential in vineyards. Journal of Applied Ecology, 1365–2664.70058. 10.1111/1365-2664.70058

Tang FHM, Lenzen M, McBratney A, Maggi F (2021) Risk of pesticide pollution at the global scale. Nature Geoscience, 14, 206–210. 10.1038/s41561-021-00712-5

Thierion V, Vincent A, Valero S (2022) Theia OSO Land Cover Map 2020. 10.5281/zenodo.6538861

Thimmegowda GG, Mullen S, Sottilare K, Sharma A, Mohanta SS, Brockmann A, Dhandapany PS, Olsson SB (2020) A field-based quantitative analysis of sublethal effects of air pollution on pollinators. Proceedings of the National Academy of Sciences of the United States of America, 117, 20653–20661. 10.1073/pnas.2009074117

Tison L, Beaumelle L, Monceau K, Thiéry D (2024) Transfer and bioaccumulation of pesticides in terrestrial arthropods and food webs: State of knowledge and perspectives for research. Chemosphere, 357, 142036. 10.1016/j.chemosphere.2024.142036

Tison L, Franc C, Burkart L, Jactel H, Monceau K, Revel G de, Thiéry D (2023) Pesticide contamination in an intensive insect predator of honey bees. Environment International, 176, 107975. 10.1016/j.envint.2023.107975

Tison L, Hahn M-L, Holtz S, Rößner A, Greggers U, Bischoff G, Menzel R (2016) Honey Bees’ Behavior Is Impaired by Chronic Exposure to the Neonicotinoid Thiacloprid in the Field. Environmental Science & Technology, 50, 7218–7227. 10.1021/acs.est.6b02658

Traynor KS, Pettis JS, Tarpy DR, Mullin CA, Frazier JL, Frazier M, vanEngelsdorp D (2016) In-hive Pesticide Exposome: Assessing risks to migratory honey bees from in-hive pesticide contamination in the Eastern United States. Scientific Reports, 6, 33207. 10.1038/srep33207

Vischetti C, Cardinali A, Monaci E, Nicelli M, Ferrari F, Trevisan M, Capri E (2008) Measures to reduce pesticide spray drift in a small aquatic ecosystem in vineyard estate. Science of The Total Environment, 389, 497–502. 10.1016/j.scitotenv.2007.09.019

Wan N-F, Fu L, Dainese M, Kiær LP, Hu Y-Q, Xin F, Goulson D, Woodcock BA, Vanbergen AJ, Spurgeon DJ, Shen S, Scherber C (2025) Pesticides have negative effects on non-target organisms. Nature Communications, 16, 1360. 10.1038/s41467-025-56732-x

Xun E, Zhang Y, Zhao J, Guo J (2017) Translocation of heavy metals from soils into floral organs and rewards of *Cucurbita pepo*: Implications for plant reproductive fitness. Ecotoxicology and Environmental Safety, 145, 235–243. 10.1016/j.ecoenv.2017.07.045

Xun E, Zhang Y, Zhao J, Guo J (2018) Heavy metals in nectar modify behaviors of pollinators and nectar robbers: Consequences for plant fitness. Environmental Pollution, 242, 1166–1175. 10.1016/j.envpol.2018.07.128

Yoder JA, Nelson BW, Jajack AJ, Sammataro D (2017) Fungi and the Effects of Fungicides on the Honey Bee Colony. In: Beekeeping – From Science to Practice (eds Vreeland RH, Sammataro D), pp. 73–90. Springer International Publishing, Cham. 10.1007/978-3-319-60637-8_5

Zaitsev GA, Dubrovina OA, Shainurov RI (2020) Iron and manganese migration in “soil–plant” system in Scots pine stands in conditions of contamination by the steel plant’s emissions. Scientific Reports, 10, 11025. 10.1038/s41598-020-68114-y

Zaller JG, Kruse-Plaß M, Schlechtriemen U, Gruber E, Peer M, Nadeem I, Formayer H, Hutter H-P, Landler L (2023) Unexpected air pollutants with potential human health hazards: Nitrification inhibitors, biocides, and persistent organic substances. Science of The Total Environment, 862, 160643. 10.1016/j.scitotenv.2022.160643

