## Supplementary Material for "Vineyard footprint revealed by honey bees used as sentinels for organic and inorganic contaminants in contrasted environments"

### Supporting Information

**Table S1 -** Internal Standards used, Retention Time, LOD/LOQ and repeatability values for synthetic pesticides residues

| | ISTD | RT | LOD ng/g | LOQ ng/g | Repeatability (RSD %) | Recovery (in % ± RSD) |
|---|---|---|---|---|---|---|
| **Fungicides** | | | | | | |
| Ametoctradin | a | 15.2 | 2.8 | 9.3 | 14.3 | 99 ± 30 |
| Azoxystrobin | a | 14.8 | 0.3 | 1.0 | 5.2 | 97 ± 22 |
| Boscalid | a | 15.0 | 2.7 | 8.9 | 14.8 | 96 ± 28 |
| Carbendazim | b | 9.8 | 0.3 | 1.0 | 4.6 | 102 ± 11 |
| Cyflufenamid | a | 16.9 | 1.8 | 6.1 | 4.9 | 99 ± 24 |
| Cyproconazole | c | 14.4 | 0.6 | 2.0 | 6.6 | 98 ± 13 |
| Cyprodinil | a | 15.9 | 7.9 | 26.4 | 15.1 | 98 ± 23 |
| Diéthofencarb | a | 14.6 | 0.2 | 0.6 | 6.4 | 96 ± 26 |
| Difenoconazole | c | 16.1 | 3.2 | 10.8 | 11.8 | 101 ± 22 |
| Dimethomorph* | a | 14.2 | 0.3 | 0.9 | 8.1 | 94 ± 21 |
| Fenbuconazole | c | 15.3 | 1.5 | 5.1 | 13.3 | 98 ± 12 |
| Fenhexamid | a | 15.0 | 4.1 | 13.5 | 8.2 | 99 ± 27 |
| Fludioxonil | d | 14.5 | 0.7 | 2.4 | 3.4 | 102 ± 10 |
| Fluopicolide | a | 15.1 | 0.7 | 2.4 | 8.4 | 100 ± 26 |
| Fluopyram | a | 15.2 | 0.3 | 1.0 | 9.4 | 97 ± 24 |
| Flusilazole | c | 15.2 | 1.8 | 5.9 | 13.3 | 100 ± 13 |
| Hexaconazole | c | 15.4 | 1.7 | 5.6 | 11.5 | 98 ± 12 |
| Imazalil | a | 13.2 | 7.5 | 24.9 | 20.7 | 99 ± 26 |
| Iprovalicarb* | a | 14.6 | 0.1 | 0.3 | 5.6 | 97 ± 17 |
| Mandipropamid | a | 14.9 | 0.3 | 0.9 | 4.1 | 97 ± 17 |
| Metalaxyl | a | 13.2 | 0.3 | 0.9 | 5.5 | 96 ± 24 |
| Metrafenone | a | 17.1 | 0.7 | 2.6 | 7.1 | 104 ± 20 |
| Prochloraz | a | 15.8 | 1.9 | 6.4 | 8.5 | 100 ± 17 |
| Pyraclostrobin | a | 16.5 | 0.2 | 0.7 | 4.5 | 101 ± 11 |
| Pyrimethanil | a | 14.4 | 5.3 | 17.6 | 10.6 | 98 ± 19 |
| Spiroxamine* | a | 13.7 | 1.2 | 4.1 | 19.5 | 99 ± 16 |
| Tebuconazole | c | 15.2 | 1.1 | 3.5 | 6.7 | 100 ± 10 |
| Thiophanate-methyl | a | 12.2 | 1.5 | 5.0 | 10.5 | 96 ± 40 |
| Triadimenol* | c | 14.1 | 1.0 | 3.0 | 5.0 | 97 ± 14 |
| Trifloxystrobin | a | 17.0 | 0.3 | 1.0 | 7.4 | 99 ± 13 |
| Zoxamide | a | 16.4 | 1.0 | 3.3 | 5.8 | 97 ± 19 |
| **Insecticides / Acaricides** | | | | | | |
| DMF (amitraz metabolite) | | 11,4 | 3.1 | 10.4 | 12.2 | 100 ± 25 |
| Acetamiprid | a | 10.5 | 0.6 | 1.9 | 6.3 | 93 ± 29 |
| Buprofezin | a | 18.2 | 0.3 | 1.0 | 13.8 | 96 ± 23 |
| Chlorantraniliprole | a | 14.0 | 3.5 | 11.6 | 7.2 | 93 ± 26 |

| | | | | | | |
|---|---|---|---|---|---|---|
| Clofentezine | a | 16.6 | 11.0 | 36.8 | 19.8 | 99 ± 27 |
| Clothianidin | a | 9.9 | 1.5 | 5.1 | 6.3 | 94 ± 31 |
| Coumaphos | a | 16.5 | 2.7 | 8.9 | 5.2 | 98 ± 17 |
| Cymiazol | a | 11.3 | 2.0 | 6.8 | 8.4 | 98 ± 34 |
| Dimethoate | a | 10.4 | 0.2 | 0.8 | 5.4 | 94 ± 25 |
| Fipronil | a | 15.9 | 1.3 | 4.2 | 6.7 | 99 ± 13 |
| Imidacloprid | a | 10.2 | 1.2 | 4.0 | 6.7 | 94 ± 29 |
| Malathion | a | 15.4 | 0.3 | 1.0 | 5.3 | 97 ± 22 |
| Methiocarb | a | 14.4 | 0.3 | 0.9 | 5.7 | 97 ± 23 |
| Methoxyfenozide | e | 15.2 | 2.4 | 8.2 | 11.4 | 100 ± 19 |
| Tebufenozide | e | 15.8 | 8.5 | 28.3 | 20.6 | 99 ± 19 |
| Thiacloprid | a | 11.3 | 1.7 | 5.8 | 4.6 | 95 ± 28 |
| Thiametoxam | a | 9.3 | 0.6 | 2.1 | 4.8 | 95 ± 26 |
| Thiodicarb | a | 12.4 | 0.2 | 0.6 | 5.5 | 96 ± 29 |
| **Synergist** | | | | | | |
| Piperonyl butoxid | a | 17.5 | 0.3 | 0.9 | 6.8 | 99 ± 14 |
| **Herbicides** | | | | | | |
| Atrazine | a | 13.1 | 1.3 | 4.5 | 7.2 | 97 ± 17 |
| Desethyl-atrazine | a | 10.2 | 1.9 | 6.4 | 13.8 | 97 ± 41 |
| Diuron | a | 13.2 | 0.5 | 1.6 | 6.0 | 97 ± 27 |
| Linuron | a | 14.5 | 3.9 | 13.0 | 11.3 | 96 ± 30 |
| Metolachlor | a | 15.6 | 0.3 | 1.0 | 6.1 | 96 ± 21 |
| Terbuthylazine | a | 14.4 | 1.6 | 5.3 | 10.9 | 98 ± 13 |
| Desethyl-terbuthylazine | a | 12.1 | 1.1 | 3.6 | 4.9 | 98 ± 17 |

*: mixture of isomers
Internal standards: a, triphenylphosphate; b, carbendazim-d4; c, tebuconazole-d6, d, fludioxonil-13C2, e, tebufenozide-d9
LOD/LOQ: concentrations corresponding to S/N ratio of 3 and 10 respectively
Repeatability: mean of the relative standard deviation (RSD) of 3 replicates (9 samples)
Recovery: mean recovery ± relative standard deviation (RSD) calculated from 39 samples

**Table S2** - Wave lengths used for the analysis of elements by ICP-AES (A) and Isotopes and mode used for the analysis of elements by ICP-MS (B) along with recovery rates of the elements

**A**

| Element | Wave length (nm) | Recovery rates (%) |
|---|---|---|
| Aluminium | 396.152 | 106.5 |
| Calcium | 422.673 | 103.2 |
| Copper | 327.395 | 101.1 |
| Iron | 259.940 | 104.9 |
| Sulfur | 181.972 | 104.9 |
| Magnesium | 285.213 | 103.6 |
| Manganese | 259.372 | 102.3 |
| Phosphorous | 213.618 | 104 |
| Potassium | 766.491 | 100.8 |
| Zinc | 213.857 | 100.4 |

**B**

| Element | Isotope | Mode | Recovery rates (%) |
|---|---|---|---|
| Arsenic | 75As | He/He | 103.9 |
| Cadmium | 114Cd | No gas | 103.5 |
| Cobalt | 59Co | He | 115.4 |
| Chromium | 52Cr | He | 96.1 |
| Molybdenum | 98Mo | No gas | 102.5 |
| Lead | 208Pb | No gas | 110.1 |

**Table S3** - Range, Mean and SD of pesticide residues quantified in the Vineyard location.

| | **Sampling** | **1** | **2** | **3** | **4** | **5** | **6** | **7** | **8** | **9** |
|---|---|---|---|---|---|---|---|---|---|---|
| | **Date** | **3-May** | **28-May** | **17-Jun** | **8-Jul** | **29-Jul** | **19-Aug** | **9-Sep** | **30-Sep** | **20-Oct** |
| **DMF** | range ppb | | | | | | | | | < 10.4 |
| | mean ± SD | | | | | | | | | < 10.4 |
| **ametoctradine** | range ppb | | 20.2 - 351.6 | 112 - 424.3 | | | | < 9.3 * | | |
| | mean ± SD | | 185.9 ± 234.3 | 268.2 ± 220.8 | | | | | | |
| **cyflufenamid** | range ppb | | 6.5 * | < 6.1 | | | | | | |
| | mean ± SD | | | < 6.1 | | | | | | |
| **difenoconazole** | range ppb | | 19.8 * | < 10.8 | | | | | | |
| | mean ± SD | | | < 10.8 | | | | | | |
| **dimethomorph** | range ppb | | 1.8 - 78.9 | 5.6 - 21.4 | < 0.9 - 0.9 | | | < 0.9 * | | |
| | mean ± SD | | 40.4 ± 54.5 | 13.5 ± 11.1 | < 0.9 ± 0.5 | | | | | |
| **fenbuconazole** | range ppb | | | < 5.1 * | | | | | | |
| | mean ± SD | | | | | | | | | |
| **fludioxonil** | range ppb | | | 4.9 -24.0 | | | | < 2.4 * | < 2.4 | |
| | mean ± SD | | | 14.4 ± 13.5 | | | | | < 2.4 | |
| **fluopicolide** | range ppb | | < 2.4 - 8.7 | 22.2 - 117.2 | | | | < 2.4 * | | |
| | mean ± SD | | 4.7 ± 5.6 | 69.7 ± 67.2 | | | | | | |
| **fluopyram** | range ppb | | | 1.1 - 3.4 | | | | | | |
| | mean ± SD | | | 2.3 ± 1.6 | | | | | | |
| **mandipropamid** | range ppb | | < 0.9 * | 5.1 - 21.1 | | | | | | |
| | mean ± SD | | | 13.1 ± 11.3 | | | | | | |
| **metrafenone** | range ppb | | | 26.6 - 59.6 | < 2.6 | | | < 2.6 | < 2.6 * | |
| | mean ± SD | | | 43.1 ± 23.4 | < 2.6 | | | < 2.6 | | |
| **pyrimethanil** | range ppb | | | | | | | 16.8 - 176.6 | | |
| | mean ± SD | | | | | | | 96.7 ± 113.0 | | |
| **tebuconazole** | range ppb | | | < 3.5 * | | | | | | |
| | mean ± SD | | | | | | | | | |
| **trifloxystrobin** | range ppb | | < 0.6 * | 7.2 - 18.7 | | | | | | |
| | mean ± SD | | | 13.0 ± 8.1 | | | | | | |
| **zoxamide** | range ppb | | | 20.1 - 109.4 | < 3.3 - 5.6 | | | | | |
| | mean ± SD | | | 64.7 ± 63.1 | 3.3 ± 3.2 | | | | | |

Values are mean of the 3 replicates in each hive (range ppb) and the mean of the two hives (mean ± SD)
* Detected or quantified in only one hive

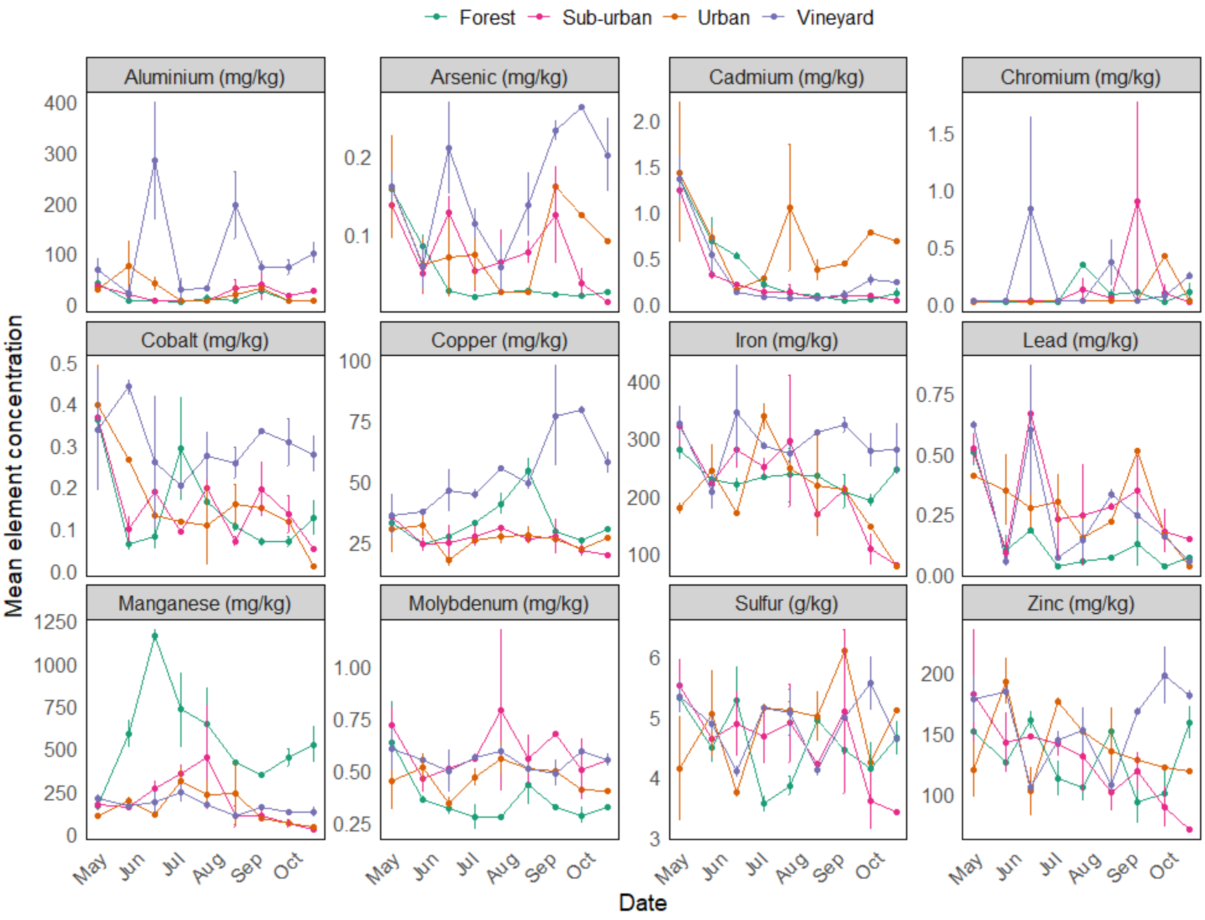


**Figure S1 -** Concentrations of elements quantified in honey bees sampled between May and October in contrasted locations. Values presented are mean ± SE of the two hives placed at each location (Forest, Sub-urban, Urban, Vineyard).

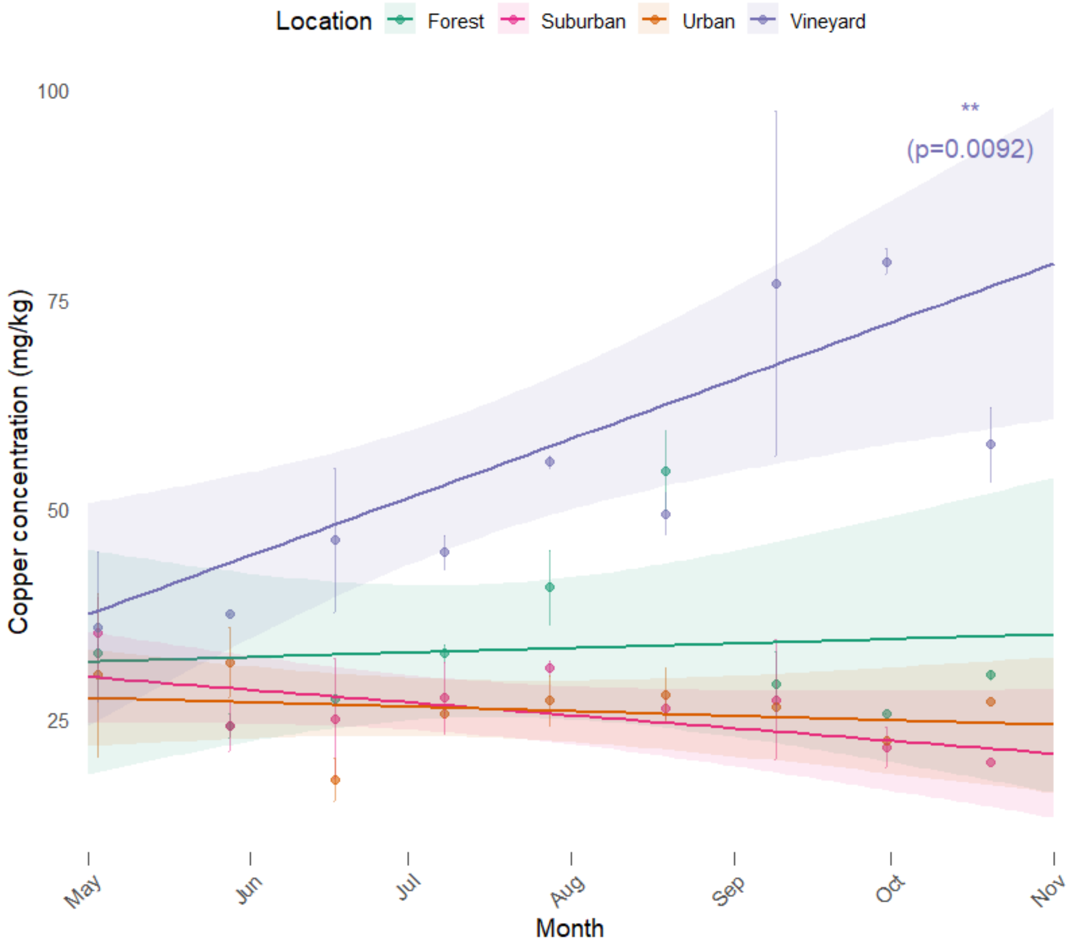


**Figure S2 -** Evolution of copper concentrations in honey bees by location. Dots are true sampling dates and fitted lines represent trends over the study period. The monthly bioaccumulation of copper was calculated through the estimation of slopes in each Location. Increasing concentrations of copper with time was revealed in Vineyards only (lm, estimate: 6.95 ± 1.95, p-value = 0.00918) whereas other locations did not exhibit any significant positive or negative relationship (p-value, Forest = 0.788, Sub-urban = 0.0986, Urban = 0.550).

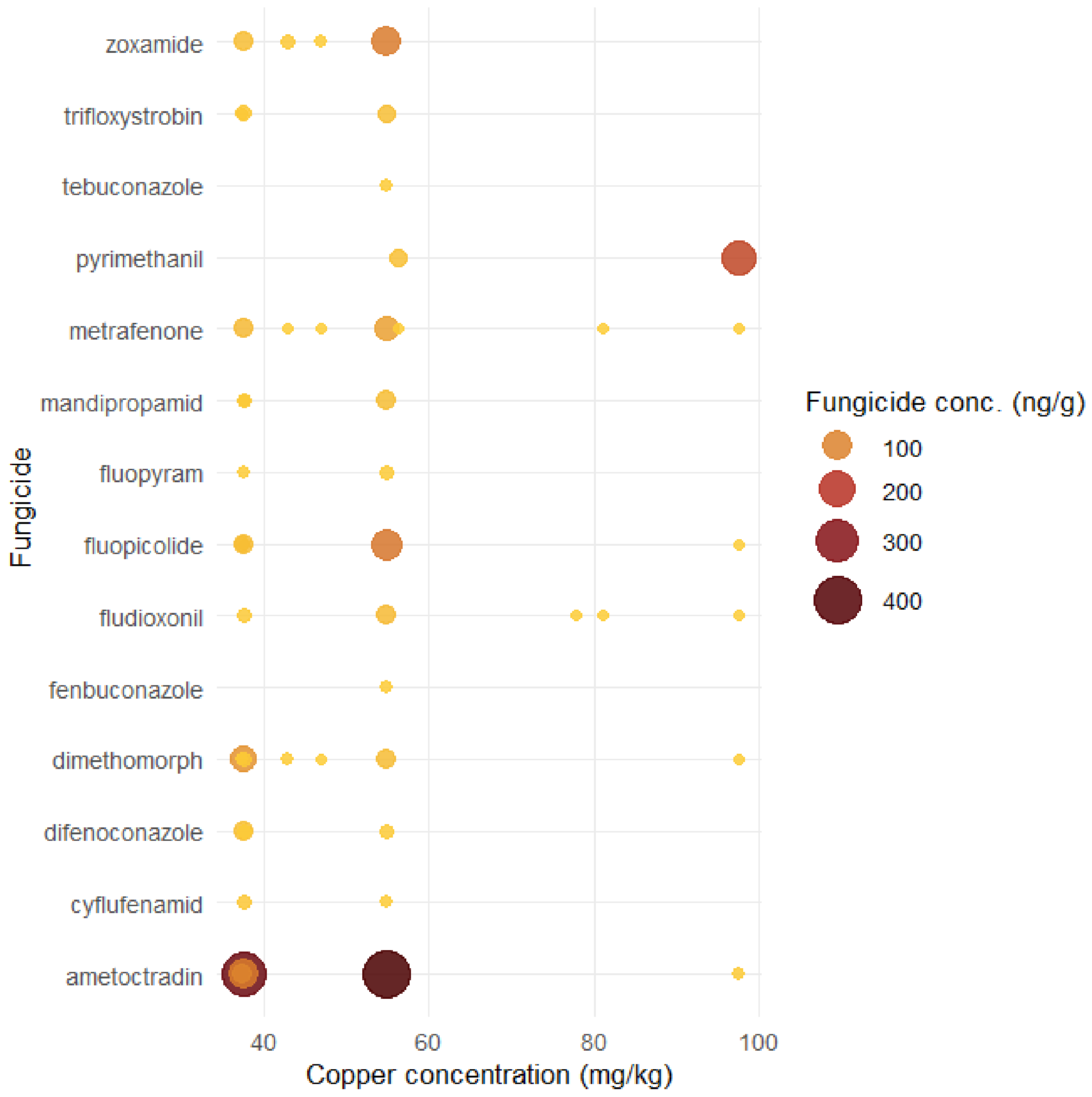


**Figure S3** - Co-occurrence of copper and fungicide residues in honeybees sampled from vineyard hives. Each point represents an sampling in which the fungicide was detected (concentration > 0). Point size and color indicate fungicide concentration (ng/g). Only samples from vineyard hives are shown. Copper concentrations are expressed in mg/kg dry weight. Spearman correlations between copper concentration and fungicide richness across samples were not significant.

**Table S4** – Acute LD50 oral and contact and RTLs of pesticide residues quantified in the Vineyard location.

| Compound | Contact acute LD50 (µg/bee) | Oral acute LD50 (µg/bee) | Pollinators RTLs |
|---|---|---|---|
| Ametoctradin | >100 | >111.5 | 2 |
| Azoxystrobin | >200 | >25 | 0.5 |
| Cyflufenamid | >100 | >100 | 2 |
| Difenoconazole | >100 | >177 | 2 |
| Dimethomorph | >102 | >32.4 | 0.648 |
| Fenbuconazole | ≥5.5 | ≥5.2 | 0.104 |
| Fludioxonil | >100 | >100 | 2 |
| Fluopicolide | >100 | >241 | 2 |
| Fluopyram | >100 | >102.3 | 2 |
| Mandipropamid | >200 | >200 | 4 |
| Metrafenone | >100 | >114 | 2 |
| Pyrimethanil | >100 | >100 | 2 |
| Tebuconazole | >200 | >83.05 | 1.661 |
| Trifloxystrobin | >200 | >200 | 4 |
| Zoxamide | >100 | >147 | 2 |

Data were collected on the PPDB database (Lewis et al., 2016)
The Lethal Dose 50 ($LD_{50}$) is the amount of a substance estimated to kill 50% of a test population when administered once (acute) by ingestion (oral) or topically (contact) within a short observation period (usually 24–48 hours)
Regulatory Threshold Levels (RTLs) are derived from acute and chronic ecotoxicity endpoint values for each substance for each species group. To account for uncertainty the endpoint values are divided by an assessment factor which follows the Tier-1 risk assessment principles for pesticide registration in the EU (see Bub et al., 2023). The most sensitive endpoint after application of the factor is the value that is used as the RTL.

**References**


Bub S, Wolfram J, Petschick LL, Stehle S, Schulz R (2023) Trends of Total Applied Pesticide Toxicity in German Agriculture. *Environmental Science & Technology*, **57**, 852–861. https://doi.org/10.1021/acs.est.2c07251

Lewis KA, Tzilivakis J, Warner DJ, Green A (2016) An international database for pesticide risk assessments and management. *Human and Ecological Risk Assessment: An International Journal*, **22**, 1050–1064. https://doi.org/10.1080/10807039.2015.1133242
